# A broadly protective CHO cell expressed recombinant spike protein subunit vaccine (IMT-CVAX) against SARS-CoV-2

**DOI:** 10.1101/2023.04.03.534161

**Authors:** Jitender, B. Vikram Kumar, Sneha Singh, Geetika Verma, Reetesh Kumar, Pranaya M. Mishra, Sahil Kumar, Santhosh K. Nagaraj, Joydeep Nag, Christy M. Joy, Bhushan Nikam, Dharmendra Singh, Pooja, Nidhi Kalidas, Shubham Singh, Mumtaz, Ashwani K. Bhardwaj, Dhananjay S. Mankotia, Rajesh P. Ringe, Nimesh Gupta, Shashank Tripathi, Ravi P.N. Mishra

**Author notes:** Dept. of Immunology and Theranostics, Center for Theranostic Studies, Beckman Research Institute, City of Hope, Duarte, California, USA. School of Medicine and Health Sciences University of North Dakota, Grand forks, ND, USA. Institute of Applied Sciences & Humanities, GLA University, Mathura, UP, India. Department of Biology, University of Marburg, 35043, Marburg, Germany (Nidhi Kalidas). ***These authors contributed equally*** (Jitender, B. Vikram Kumar, Sneha Singh).

## Abstract

Protective immunity induced by COVID-19 vaccines is mediated mainly by spike (S) protein of severe acute respiratory syndrome coronavirus 2 (SARS-CoV-2). Here, we report the development of a recombinant prefusion stabilized SARS-CoV-2 spike protein-subunit-based COVID-19 vaccine produced in the mammalian cell line. The gene encoding ectodomain (ECD) of the spike protein was engineered and cloned into Freedom pCHO 1.0, a mammalian expression vector, and subsequently expressed in the Chinese Hamster Ovary suspension cell line (CHO-S). The recombinant S protein ectodomain (hereafter referred to as IMT-CVAX) was purified using a combination of tangential flow filtration and liquid chromatography. Biochemical and biophysical characterization of IMT-CVAX was done to ensure its vital quality attributes. Intramuscular immunization of mice with two doses of adjuvanted IMT-CVAX elicited a strong anti-Spike IgG response. In pseudovirus-based assays, IMT-CVAX– immune mice sera exhibited a broad-spectrum neutralization of several SARS-CoV-2 variants of concern (VoCs). Golden Syrian Hamster immunized with IMT-CVAX provided excellent protection against SARS-CoV-2 infection, and, hamster immune sera neutralized the live SARS-CoV-2 virus. The adjuvanted IMT-CVAX induced robust T_fh_-cells response and germinal center (GC) reaction in human ACE2 receptor-expressing transgenic mice. The findings of this study may pave the way for developing next-generation protein subunit-based vaccines to combat the existing SARS-CoV-2 and its emerging VoCs. The IMT-CVAX is produced using a scalable process and can be used for large-scale vaccine production in an industrial setup.

## Introduction

The COVID-19 pandemic caused by the Novel Coronavirus SARS-CoV-2 has had a catastrophic effect on the world’s population resulting in more than 760 million infections and 6.8 million deaths worldwide (1). Because of its unprecedented global spread, morbidity and mortality, the World Health Organization (WHO) declared it a pandemic in March 2020 (2).

Currently, several COVID-19 vaccines are licensed and rolled out for human use the world over. These vaccines are produced using various platforms such as mRNA (Moderna, Pfizer), whole inactivated virus (Bharat Biotech, SinoPharm, Sinovac, Valneva), viral vector (AstraZeneca, J&J, CanSino), DNA (Zydus Cadila) and recombinant protein subunit (Novavax, Biological E. Ltd., Sanofi & GSK) vaccines etc (3). Most licensed COVID-19 vaccines target Spike protein, emphasizing its relevance in viral pathogenesis (2–5). To tackle the emergence of differentially infective variants, the concomitant rise in vaccine breakthrough infections, the waning of vaccine-induced immunity, and the assurance of continuous global supply and comprehensive coverage, large quantities of COVID-19 vaccines must be produced using different technologies.

Protein subunit-based vaccines represent an attractive class of vaccines with a remarkable history boasting of their safety and efficacy. Protein-based vaccines have been effective against a diverse range of infections and can be administered in almost all age groups, from neonates to the elderly (6, 7). Another essential aspect of protein-based vaccines is the ease of large-scale production, storage in standard refrigerators, and affordability. When it comes to combating a fast-replicating virus with high mutating potential, such as SARS-CoV-2, protein subunit-based vaccines may prove advantageous for the reasons mentioned above (8).

The SARS-CoV-2 is an enveloped virus with a positive sense, single-stranded RNA of nearly 30,000 nucleotides that encode four structural proteins: spike (S), membrane (M), envelope (E), and nucleocapsid (N). The S protein, a surface-exposed, homotrimeric glycoprotein, is the major virulence factor and antigenic determinant of SARS-CoV-2 (9, 10). It decorates the viral surface and interacts with the host cell receptor ACE2 (Angiotensin Converting Enzyme 2) resulting in host-viral membrane fusion and entry of the virus inside the host cell (11–14). The trimeric Spike protein is made of three glycosylated monomer units, each with a molecular weight of 180-200kDa. Each monomer consists of an extracellular N-terminus, a transmembrane (TM) domain anchored in the viral membrane and a short intracellular C-terminal segment (15).

The S protein of SARS-CoV-2 belongs to the family of class-I fusion proteins, which are known for their innate instability making it difficult to express them in their native forms. The S protein exists in two distinct conformational states: i) prefusion, antigenically intact but metastable and, ii) postfusion, highly stable but antigenically compromised (16–19). Therefore, to be able to use the intact S protein or its ectodomain (ECD) as a vaccine antigen, stabilization of its prefusion conformation is required. It has been previously reported that prefusion-stabilized protein exhibits better expression in recombinant hosts and is much more stable than its wild-type counterpart (17).

We followed a combination of different protein engineering approaches as reported in the recent study, for designing the prefusion stabilized S protein ECD-based vaccine antigen, including substitutions of certain amino acids with proline, which increases the loop stability, modification of the furin cleavage site and insertion of a trimerization motif in the protein (17). In addition, we replaced the native secretion signal sequence with a heterologous signal sequence for better secretion of protein, and removed the transmembrane domain from the C-terminus of the protein. The designed gene construct was then exogenously expressed by using a combination of cGMP (current good manufacturing practices)-banked mammalian cell line (CHO-S) and expression vector (pCHO1.0) which permits achieving a high yield in a high-density cell culture system. Noteworthy, the feasibility and potential of this expression system have already been demonstrated in our recent research for the expression of industry-compatible recombinant monoclonal antibody (20).

Adjuvants play a pivotal role in the development of viral and bacterial vaccines. Adjuvants are capable of enhancing the breadth of antibody and CD4^+^ T cell responses and/or shaping antigen-specific immune responses, leading to relatively better protection (21–24). Moreover, the choice of appropriate adjuvant remains a key factor for COVID-19 vaccines, where it is important to ensure a balanced humoral and cell mediated immunity (25). The rationale of using different adjuvants, such as Alum (2% Alhydrogel), AddaVax^TM^, (a squalene-based oil-in-water nano-emulsion, similar to MF59 adjuvant), and Alum + CpG ODN 2006 (synthetic immunostimulatory oligonucleotide-ODN containing unmethylated CpG dinucleotides), is to evaluate the impact of the nature of adjuvants on immunogenicity, viral neutralization and in-vivo efficacy. Alum is one of the oldest approved adjuvants used for several licensed vaccines and is primarily skewed towards Th-2 mediated immune response, whereas CpG ODN 2006 induces primarily the Th-1 response. AddaVax^TM^ is a MF59-like adjuvant. MF59 possesses the ability to induce strong antigen-specific CD4+ T-cell responses and to induce strong and long-lasting and broad memory T- and B-cell responses. It is known that MF59 effectively induces both cellular as-well-as humoral immune responses and its post-boost recall responses are quite robust, even after several years of primary immunization (26, 27).

Establishing a germinal center (GC) response is critical for developing long-term protective humoral immunity. Vaccine-induced antigen-specific B and follicular helper CD4^+^ T (T_fh_) cells response in SARS-CoV-2 vaccines is reported to be a significant contributory factor for excellent immunogenicity and high virus-neutralizing antibody titers (28–33). However, the potential of an adjuvanted COVID-19 protein subunit vaccine for the generation of germinal center B and T_fh_ cells response is poorly understood, which we seek to address in the present study.

To have a rapid response to the pandemic, it is imperative to have a robust process capable of generating high quality and quantity spike protein that meets the perquisites for fundamental and translational research and pre-clinical assessments. In this paper, we provide a proof-of-concept of a holistic process for the rapid development of a safe, efficacious and affordable COVID-19 vaccine based on a protein subunit approach. To the best of our knowledge, an integrated approach encompassing the design of antigen, the robust industrial process for tag-free spike protein expression using CHO-S suspension cell line and the proof of concept of the efficacy and viability of a vaccine candidate in three different small animal model (naïve Mice, hACE2 transgenic mice & hamster) system is reported for the first time.

Looking back at the past scenario and considering the present and future demands of COVID-19 vaccines, industry-compatible platforms and processes for producing biopharmaceutical-grade proteins are the need of the hour. If/when situations similar to the recent pandemic arise, such pre-planned endeavors may prove crucial for rapid vaccine proof of the concept development as well as for research, diagnostics and serological applications.

## Results

### Prefusion stabilized Spike protein antigen IMT-CVAX (pc DNA 3.4–IMT-C20) showed good transient expression in the ExpiCHO Cell line

Transient expression is commonly used for rapid confirmation of protein expression. ExpiCHO cells transfected with pcDNA 3.4–IMT-C20 construct was cultured for 12 days. The viability of the transfected cells remained at≥85% until Day 9, post which the viability started dropping, and therefore, the culture was harvested on Day 12 at 73.8% viability. The VCD was highest on Day 5 (∼126.5×10^5^ cells/mL) and dropped to ∼ 39.5 ×10^5^ cells/mL at the time of harvest (Fig 1b). Our previous study demonstrated that expression of spike protein in ExpiCHO cells is visible in SDS-PAGE approximately 72 hours (Day 3) post-transfection (Data not shown). The SDS-PAGE analyses of crude culture supernatant collected on days 3-11 revealed that protein expression in transient culture was detectable from Day 3 (∼72h) and showed a gradual increase till day 8. A saturation in expression was observed from Day 8 onwards (Fig 1c). A drop in VCD was observed after Day 5, but it did not have any detrimental impact on protein expression, as shown in SDS-PAGE and western blotting results (Fig 1c).

**Fig. 1:**
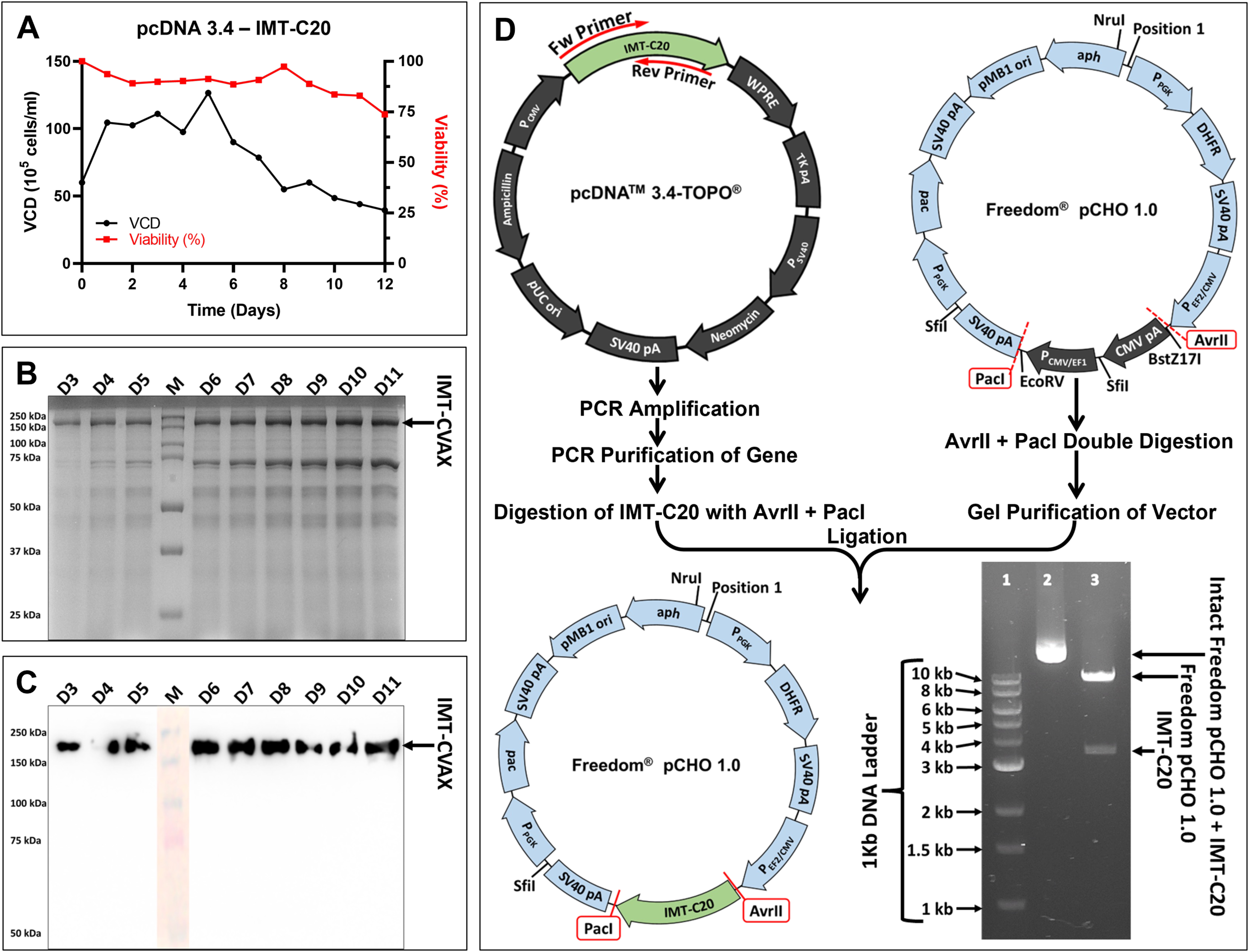
Transient expression of the pcDNA 3.4–IMT-C20 construct and cloning of IMT-C20 into Freedom pCHO 1.0. **(A)** is the viability and viable cell density (VCD) of ExpiCHO cells transfected with the pcDNA3.4–IMT–C20 construct during transient production. **(B and C)** are the SDS-PAGE and Western Blot, respectively, of the day-wise samples of transfected ExpiCHO cells from Day 3 to Day 11. **(D) IMT-C20 cloning in Freedom pCHO 1.0:** The gene (IMT-C20) was amplified from pcDNA 3.4 using gene-specific primers designed such that the forward and reverse primers had sites for AvrII and PacI restriction enzymes, respectively. The amplified gene was purified and digested using AvrII and PacI. Freedom pCHO 1.0 intact vector was also double-digested using the same enzymes (AvrII and PacI) and the larger fragment was extracted from 1% agarose gel. IMT-C20 was ligated in Freedom pCHO 1.0 and was immediately used to transform chemically competent *E. coli* DH5α cells. Freedom pCHO 1.0–IMT-C20 construct was isolated from the transformed E. coli cells and was double-digested with AvrII and PacI to confirm the ligation. Lanes 2 and 3 in the agarose gel image depict the undigested and the digested Freedom pCHO 1.0– IMT-C20 construct, respectively.

The purification process of His-Tagged IMT-CVAX was very straightforward. The clarified spent media was subjected to TFF for buffer exchange and diafiltration. Subsequently, purification of IMT-CVAX was done using either affinity chromatography alone or a combination of affinity and size-exclusion chromatography. Cultures having viabilities ≥85% at the time of harvest yielded highly pure IMT-CVAX after just one round of affinity chromatography, which was performed using the binding and elution methodology, with the bound IMT-CVAX being eluted at 100% gradient of elution buffer (SI Appendix, Fig 1a, gel-lane 3), after a thorough wash with 10% elution buffer. Desalting was done as a means of buffer exchange (SI Appendix, Fig 1a, gel-lane 5). An additional round of size-exclusion chromatography was required after the affinity chromatography for culture with viabilities <85%. There were minute quantities of some cellular proteins which could not be removed entirely during affinity chromatography (SI Appendix, Fig 1b, left gel-lane 6), but were separated during size-exclusion chromatography, yielding highly pure IMT-CVAX (SI Appendix, Fig 1b, right gel-lanes 3 and 4). Desalting was not needed after size exclusion.

### CHO-S and Freedom pCHO 1.0 were found to be a good system for achieving a high yield of IMT-CVAX

We initially expressed IMT-CVAX with histidine tag during transient expression using research-grade cell line and vector (i.e. ExpiCHO & pcDNA 3.4). However, transient expression is not a viable way to express recombinant protein for biopharmaceutical applications as it suffers from scalability limitations and industry compatibility. Therefore, to meet the regulatory and industry expectations, we have expressed the tag-free protein using cGMP banked CHO suspension cell line (CHO-S) and expression vector pCHO 1.0. With gene-specific primers, IMT-C20 was amplified from pcDNA 3.4 excluding the sequence encoding 8x histidine tag and cloned into Freedom pCHO 1.0 industry-grade vector. Freedom pCHO 1.0 is a common expression vector used for high-density cell culture-based production of therapeutic proteins. The clones of interest were confirmed by digestion of Freedom pCHO 1.0–IMT-C20 using AvrII and PacI restriction enzymes (Fig-1d). Subsequently, the recombinant pCHO 1.0–IMT-C20 vector was linearized using NruI and used for transfecting CHO-S cells. The transfected cells were propagated in CD FortiCHO medium under increasing concentrations of two selection reagents (Puromycin and Methotrexate), to facilitate the chromosomal integration of the gene for stable cell line generation. At the end of the selection phase, four stable cell pools (viz C20-1a, C20-1b, C20-2a, and C20-2b) were generated. Based on the volumetric productivity from the batch production data (*SI Appendix*-Figure 3i) at the flask level, C20-2a was selected for further optimization of feed and culture conditions.

### IMT-CVAX was highly expressed in CHO-S Fed-batch suspension cell culture

C20-2a, having the highest expression of IMT-CVAX, was chosen for further optimizations of feeding and temperature conditions in fed-batch mode. As mentioned in Tables 1 (*SI Appendix*, Table 1), feeding was done, and a temperature shift from 37℃ to 32℃ was also done in Runs 6-8 (*SI Appendix*-Table 2). Cell viability and VCD of Runs 6-8 were significantly higher than those of Runs 1-5. In the case of Runs 2 and 3, the cell viability dropped below 75% after days 9 and 10, respectively. This drop in cell viability may be due to a limitation of substrates during production (*SI Appendix*-Figure 3b, 3c). On the contrary, Run 7 (2.5%, 32℃ on day 5) had the best viability (*SI Appendix*-Figure 3g), but the expression of IMT-CVAX was relatively lower when compared to other runs (*SI Appendix*-Figure 3i). The highest expression of IMT-CVAX was observed in Run 4 (2.5% feed, 37℃) and Run 6 (5% feed, 32℃). The cell viability observed in Run 4 was marginally better than that of Run 6, whereas the VCD observed in Run 6 was much better than that of Run 4. Higher VCD at the time of harvest would result in minimal quantities of cellular proteins in the spent medium, thereby making downstream processing easier. Therefore, the culture conditions of run number six were considered for further production of IMT-CVAX in a stirred tank bioreactor.

### High yield production of IMT-CVAX in bioreactor

Following optimization of feeding strategies and culture conditions, especially temperature, the selected stable cell pool C20-2a was cultured in a bench-top stirred tank bioreactor for bulk production of IMT-CVAX. In-process samples were collected every 24 hours. The data generated following sample analyses indicated that the cell viability remained above 85% until day 13 and a slight drop in viability (below 80%) was observed after day 13. The highest VCD was observed on day 7, whereas a gradual drop in VCD was observed after day 11 (Fig.-2a). The sample collected on day 14, just before harvest, was analyzed by SDS-PAGE (neat, 1:2, and 1:4 dilution) (Fig.-2c). The yield of tag-free IMT-CVAX, as evidenced by the densitometric analyses, was ∼440 μg/mL (Fig.-2d).

### Purification of tag-free IMT-CVAX

The purification process of proteins without affinity tags comes with inherent challenges and requires rigorous downstream processing to recover high purity proteins. In the present study, a simple process for the purification of tag-free S protein was developed successfully. The purification of IMT-CVAX expressed by the stable cell pool C20-2a with viability ≥85% at harvest is a 3-step process, as depicted in the diagram (SI Appendix, Fig-5b). The harvest was first clarified by combining centrifugation and filtration using a 0.22µm filter to remove the cell mass and debris. The second step involved TFF; more minor protein impurities were filtered out and the culture medium was concentrated and exchanged with the chromatography buffer. The third step involved the chromatographic separation of IMT-CVAX from other impurities and was dependent on the viability of cells at the time of harvest. Whenever cell viability was ≥85%, a single round of either anion-exchange or size-exclusion chromatography was sufficient to obtain highly pure IMT-CVAX (Fig 2D: gel lane 2, and 2F). On the contrary, when the viability of cells at the time of harvest was <85%, a two-step chromatography involving anion exchange with CaptoQ resin as the capture step and subsequently size-exclusion chromatography as a polishing step was required to achieve the desired level of purity (Fig 2D: gel lanes 5-11, and 2E). It is clear from the experimental data that ensuring cell viability ≥85% during the production phase allows purification using only single-step chromatography.

**Fig. 2:**
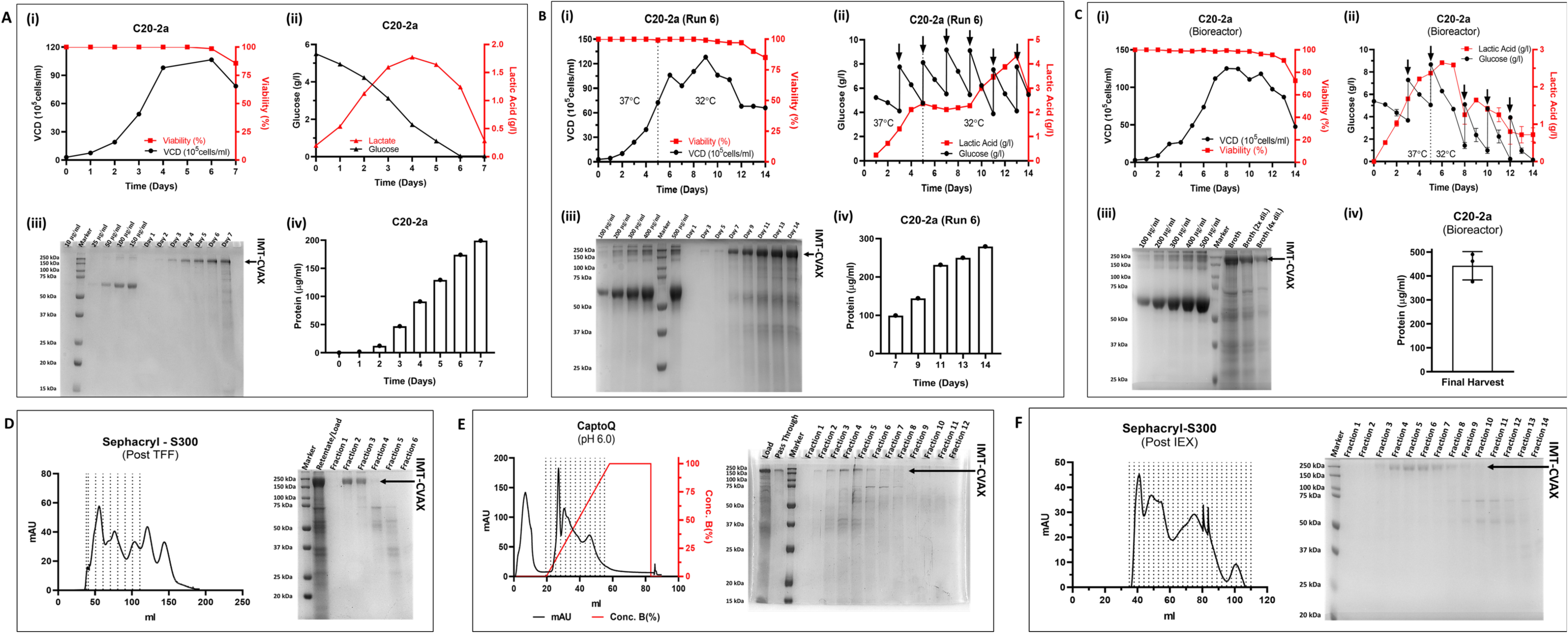
Generation of Stable Cell Pool, Optimization of Culture Conditions, Scale-up, and Purification of IMT-CVAX: (A) Stable Cell Pool with highest expression of IMT-CVAX (C20-2a)was selected for feed and temp optimizations (i) The stable pool C20-2a, at the time of harvest, had 85.7% viability and a VCD of 78.50×10^5^ cells/ml. (ii) is the glucose and lactate profile during the batch run. (iii) The daily samples collected during the batch production were resolved on 10% polyacrylamide gel under reducing conditions. Densitometric analyses of the SDS-PAGE demonstrating the protein concentration present in the daily samples. (iv) Comparing with the known concentrations of BSA (lanes 1, 3-6 in gel image), the relative yield of IMT-CVAX was ∼199.27 µg/ml. (B) Feeding and temperature conditions yielding high expression of IMT-CVAX without compromising the viability of cells were selected for scale-up: Adding 5% feed on alternate days from day 3 onwards and a 37℃-32℃ temperature shift on day 5 maintained the viability >80% till day 14 (i and ii). As per the densitometric analyses performed from the SDS gel (iv), the highest concentration of IMT-CVAX was 279.31µg/mL on Day 14 (iii). (C) Scale-up using the optimize culturing conditions: (i and ii) are the cell viability and metabolite (glucose and lactic acid) profiles. (iii) Densitometric analyses of the sample collected at the time of harvest yielded in ∼ 440µg/ml of IMT-CVAX. Known dilutions of BSA (100-500 µg/ml in lanes 1-5) were used as references (iii) for the densitometric calculation of protein yield (iv). (D) One-step chromatography: the retentate from TFF was concentrated and loaded directly onto Sephacryl-S300 size-exclusion column. Highly pure IMT-CVAX (lanes 4 and 5) was obtained. (E and F) Two-step Chromatography: (E) anion-exchange chromatography using CaptoQ with buffers of pH 6. Pure IMT-CVAX was obtained in the binding pass-through (lane 2 from the left). (F) Fractions collected during elution of anion-exchange (lanes 5-11 in gel in Panel-E) were pooled and purified using size-exclusion chromatography.

### Biophysical and biochemical Characterization of IMT-CVAX

IMT-CVAX, when run under reducing conditions in SDS-PAGE (10% gel), appeared to migrate with an observable molecular weight of ∼180 kDa, on par with the theoretical molecular weight of the S protein monomer. This finding was confirmed further by carrying out western blotting (Fig-3a) and LC/MS analyses under reduced conditions (Fig-3c).

**Fig. 3:**
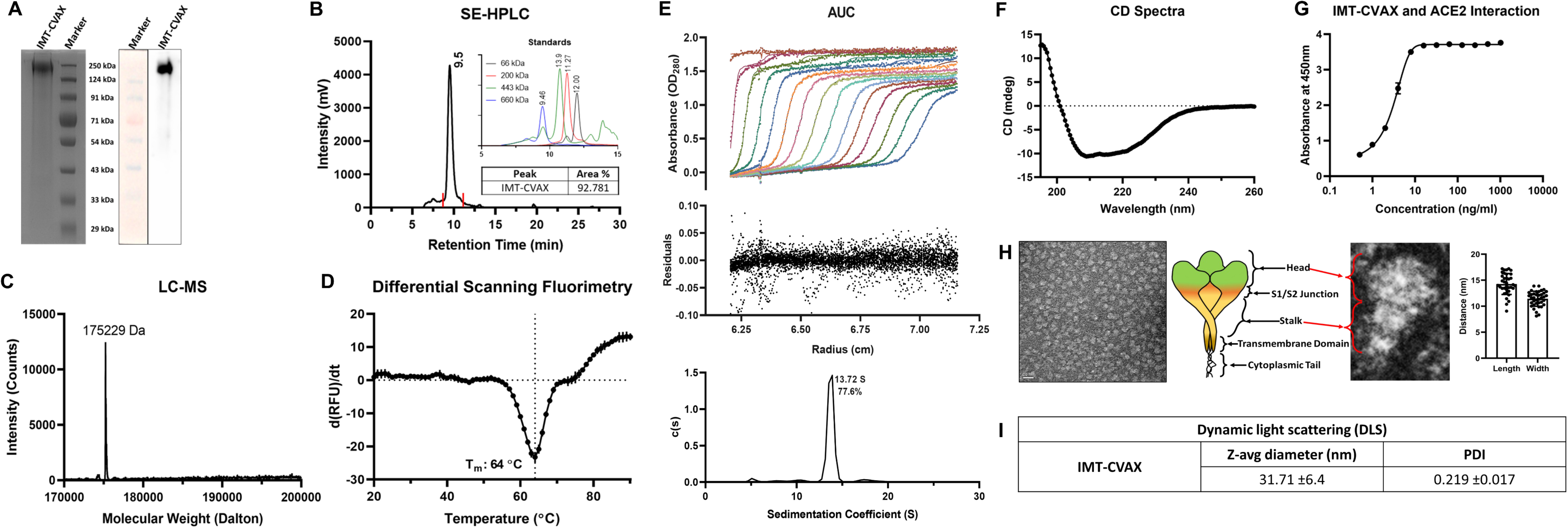
Biophysical Characterization of IMT-CVAX. **(A)** SDS Page and western blot of pure, untagged IMT-CVAX. **(B)** SE-HPLC chromatogram showing one major peak of IMT-CVAX at 9.5 ml, occupying >92% of pure protein (area% mentioned in the inset table in the bottom right corner). The approximate molecular weight of IMT-CVAX is close to the 660 kDa molecular weight standard (as depicted in the inset at the top-right corner). The chromatogram also shows the presence of a small fraction of larger aggregates (∼ 5% of total area) of IMT-CVAX. **(C)** The molecular weight of reduced IMT-CVAX, as depicted in the LC-MS spectrum, is 175.229 kDa. **(D)** The thermal denaturation of IMT-CVAX between 20℃-95℃ with a ramp rate of 0.5℃/min, revealed that its melting temperature is 64℃. **(E)** IMT-CVAX (in 20mM Tris, 200mM NaCl, pH 8.00) was subjected to SV AUC and absorbance at 280nm was measured at an interval of 3 min and fitted to a continuous c(s) distribution from 0 S to 20 S. Only 15 alternate scans are shown here for clarity. Upper panel shows A_280_, middle panel shows the residuals of the fitted data, and lower panel shows the c(s) distributions (Absorbance, residuals and c(s) distribution plot). **(F)** Far-UV CD spectrum of IMT-CVAX. The lowest negative value corresponding to 209nm suggests abundance of β-sheets. **(G)** Graph showing the interaction of IMT-CVAX with ACE2. **(H)** Images obtained from nsEM show the trimeric molecules of IMT-CVAX. Images of 40 individual molecules from 4 nsEM images were measured. One trimeric molecule of IMT-CVAX is approximately 14nm long and 11nm wide. The different domains of spike are co-related with the cartoon image. **(I)** Z-average diameter and polydispersity index obtained from dynamic light scattering analyses.

To explore the formation of trimer SE-HPLC, analyses of IMT-CVAX were performed under non-reducing conditions. It was observed that IMT-CVAX appeared to have a retention time corresponding to ∼660 kDa, the approximate molecular weight of the S trimer (Fig 3b). Furthermore, a fraction of aggregates (∼ 7%) was also observed in SE-HPLC analyses. This is imperative to mention that similar data was also reported in a previous study (34).

The Z-average diameter (Z-avg) and polydispersity index (PDI) deduced using DLS analyses were 31.71 + 6.4 and 0.219 + 0.017, respectively, indicating that the IMT-CVAX had uniformity in terms of shape, size, and mass (Fig-3i). In addition, the AUC analyses demonstrated that the significant proportion of the IMT-CVAX in solution (77.6%) was uniform. However, a small fraction with sedimentation coefficients higher than those of individual molecules of IMT-CVAX was also observed, indicating that the IMT-CVAX had a slight propensity for aggregation (Fig-3e) in the present form. IMT-CVAX showed a sedimentation coefficient of 13.72 based on the c(s) distribution. The overall data corroborates with the data obtained from SE-HPLC. Thermal denaturation assays revealed that the IMT-CVAX has a single melting temperature (Tm) of ∼64℃ (Fig 3-D).

The confirmation of secondary structures in the IMT-CVAX was determined by using CD spectroscopy. As depicted in Fig-3f, the single, lowest negative value corresponding to 209 nm wavelength indicated the prominence of β-sheets. When analyzed using the BeStSel webserver, the acquired data indicated the prominence of secondary structures. In addition, we attempted to elucidate pure IMT-CVAX using negative stain transmission electron microscopy (nsTEM). We observed that IMT-CVAX is present in its trimeric form. The data confirms that the IMT-CVAX has a ‘clove-like’ head and stalk structure corresponding to the S1 and S2 domains. The average lengths and widths of individual molecules were ∼14 nm and ∼11nm, respectively (Fig-3h). Similar data reported by another group substantiated well with our findings (34).

### IMT-CVAX showed a high binding affinity for human ACE2 receptor

The affinity of the interaction between the IMT-CVAX and human ACE2 receptor was determined by ELISA. IMT-CVAX, serially diluted, was allowed to interact with human ACE2 receptor coated on an ELISA plate and subsequently detected using anti-His antibody–HRP conjugate. The results indicated a high affinity for the interaction between IMT-CVAX and the hACE2 receptor (Fig-3g).

### Adjuvanted IMT-CVAX elicits potent Anti-Spike IgG antibodies in BALB/c Mice

Immunogenicity of various formulations of the adjuvanted (AddaVax, Alum, and Alum + CpG ODN 2006) IMT-CVAX was assessed in BALB/c mice. Anti-Spike IgG was quantified by indirect ELISA of the serum samples collected from immunized mice (Fig-4a). An excellent anti-Spike IgG response was generated in mice against IMT-CVAX formulated with different adjuvants (Fig-4b).

### Anti–IMT-CVAX IgGs in mice sera effectively neutralize different Variants-of-Concern (VoCs) of SARS-CoV-2

The mouse serum after IMT-CVAX immunization was evaluated for neutralization of different variants: Alpha, Beta, and Delta, along with Wild Type (D614G) of SARS CoV-2 in a pseudovirus-based neutralization assay. Although IMT-CVAX is made in the background of the Wild-Type sequence, the sera exhibited effective cross-neutralization of other variants. Cross-neutralization is a critical characteristic of an effective vaccine candidate against rapidly mutating viruses. Noteworthy, the mice sera tested had a relatively lower neutralizing antibody titer against the Delta variant compared to the other pseudotypes chosen for the test. There was no vaccine formulation-related difference in protection breadth and magnitude of neutralizing antibodies. However, a marginal betterment was observed with adjuvants AddaVax™ and CPG 2006 (Fig-4c).

### IMT-CVAX vaccination confers excellent protection against SARS-CoV-2 infection in Hamsters

Hamsters are an excellent small animal model to study disease pathogenesis and vaccine development for SARS-CoV-2 because of their innate susceptibility to the virus and ability to recapitulate many of the respiratory distress symptoms that infected humans exhibit. In this study, we used 5-6 weeks-old Golden Syrian Hamsters to assess the pre-clinical efficacy of the IMT-CVAX as a potent COVID-19 vaccine. Briefly, the hamsters were divided into three groups: control uninfected, control infected, and vaccinated. The vaccinated groups received three doses of IMT-CVAX adjuvanted with AddaVax in a ratio of 1:1 at an interval of 14 days. Two weeks post-dose 3, the hamsters were challenged with SARS-CoV-2 (Isolate Hong Kong/VM20001061/2020, Cat No.: NR-52282, BEI Resources, NIAID, NIH) and were closely monitored for the next 14 days. Studying the physical and anatomical clinical signs post-challenge with SARS-CoV-2 and comparing them with the control groups, we found that the groups administered with IMT-CVAX were completely protected against SARS-CoV-2 infection. The physical clinical signs of viral challenge manifested as loss of body weight, lethargy, piloerection, abdominal respiration, and hunched back. The infected control animals showed maximum clinical signs up to day 4 post-infection (p.i.), after which a gradual recovery was observed until day 14 p.i. Not only did the IMT-CVAX + AddaVax group exhibit significantly reduced clinical signs compared to the infected controls, but its recovery phase also started (by day 3) earlier than the infected control group (Cite appropriate figure number). A steady decline in the total body weight of the control infected hamsters were observed up to day 2 p.i., compared to uninfected animals. Though not entirely, a slight recovery in body weight was observed till day 4 p.i. A good rescue in body weight was observed in the case of hamsters administered with the IMT-CVAX + AddaVax formulation when compared to the other groups. Also, by day 14 p.i., all the animals in both control infected and vaccinated group had regained their lost body weight and was comparable to the uninfected control group. The group vaccinated with IMT-CVAX + AddaVax recovered faster than infected control animals, starting from 2 days p.i.

Compared to the other groups on day 4 p.i., the lungs from the control infected animals showed moderate gross lesions among both males and females. The hamster lungs from the IMT-CVAX + AddaVax group showed only mild lesions among male and female hamsters. On day 14 p.i., control infected female hamsters showed mild lung lesions, but no such lesions were observed among male and female hamsters in the IMT-CVAX + AddaVax group. (Fig. 5g).

**Fig. 5:**
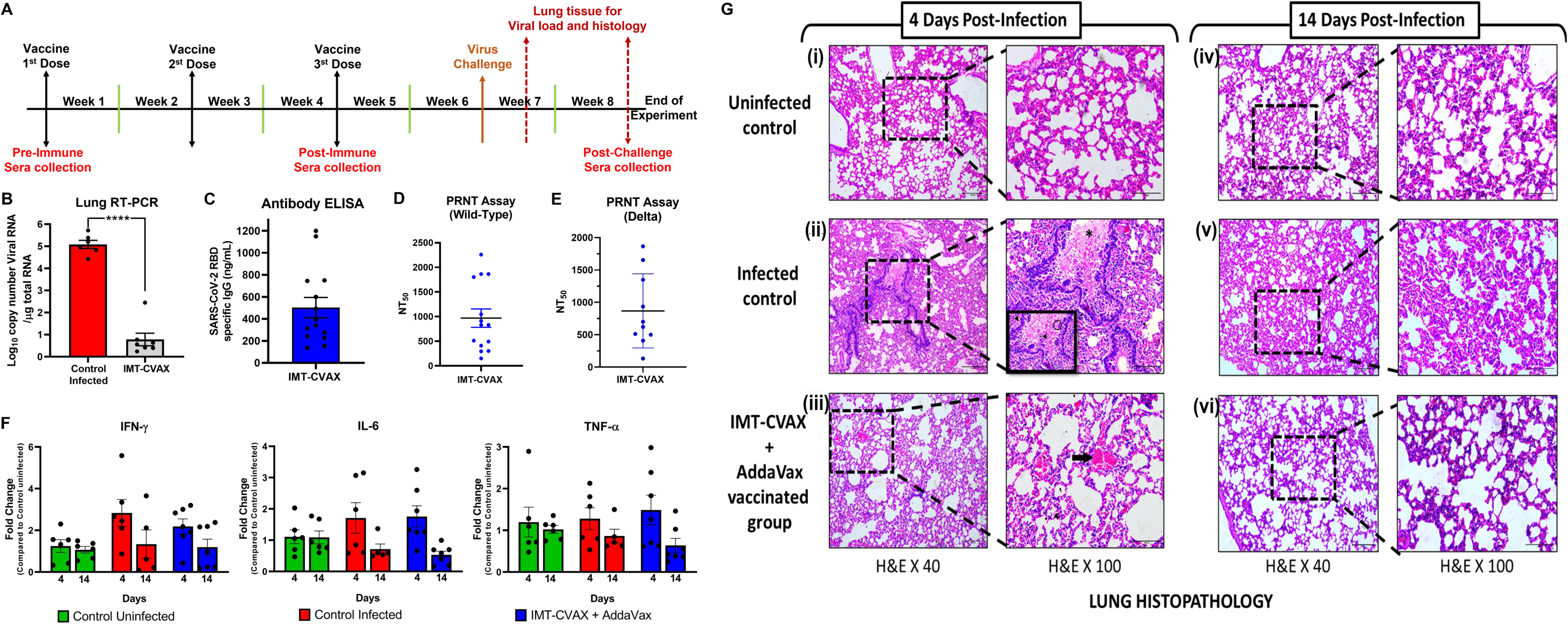
Pre-Clinical Efficacy of IMT-CVAX in Hamsters. (A) Quantification of lung viral RNA load post-infection. Lungs were harvested (Day 4 p.i.) and used for estimation of viral load by qRT-PCR. The viral RNA copy number was estimated using a standard curve generated from Ct values. IMT-CVAX + AddaVax treated hamsters had a significant reduction (∼ 5 times) in lung viral load when compared to the control infected group. **(B)** PRNT Assay: The sera from vaccinated Syrian Golden Hamsters pre-SARS-CoV-2 (HK strain) challenge infection was used to find out the 50% neutralization titer against SARS-CoV-2 HK strain. The data were represented as NT50. **(C)** Anti-spike RBD IgG ELISA. The sera from vaccinated Syrian Golden Hamsters before SARS-CoV-2 (HK strain) challenge infection was used to quantify the anti-SARS-CoV-2 IgG specific to spike RBD protein using an IgG standard plot. **(D)** Lung cytokine mRNA level: The IFN-ϒ/IL-6/TNF-α cytokine mRNA level was estimated by qRT-PCR from total RNA isolated from the lungs of control/vaccinated Syrian Golden Hamsters post-SARS-CoV-2 (HK strain) challenge infection from day-4 and day-14 group. The data were normalized with 18s RNA and represented as fold change in expression relative to control. **(E)** Lung histopathology. The lung from control/vaccinated Syrian Golden Hamsters on Day 4 and Day 14 (Hongkong strain) challenge infection was examined microscopically based on 3 histological criteria and scored. The data were represented as mean ± SEM. (i) Lung sections showing normal alveolar histology. (ii) Bronchiole lumen filled with exudates(asterisk), denuded necrotised epithelial cells (circle) as well as neutrophil and mononuclear cell infiltration (arrowheads) and diffuse alveolar septal thickening. (iii) Congested blood vessels (arrow). (iv) Lung sections showing normal alveolar histology. (v) Moderate alveolar septal thickening with inflammatory cell infiltration. (v) Mildly thickened alveolar walls.

Total RNA isolated from the lung tissues obtained from the control infected group had approximately 5-fold higher viral RNA when compared to the group immunized with AddaVax adjuvanted IMT-CVAX at day 4 p.i. However, it is imperative to note that the levels of viral RNA eventually receded in the control infected group and were comparable to the vaccinated group by day 14 p.i. owing to the natural immune response mounted in the animals against the virus (Fig. 5bi). Quantification of the expression of cytokine mRNA in lung tissue by qRT-PCR revealed that the levels of IFN-γ were significantly elevated in the control infected group than in vaccinated groups at day 4 p.i. with no apparent difference in the levels observed at day 14 p.i. This finding indicates the persistence of viral infection in the lungs from control infected animals, which correlates with their viral load. Levels of cytokines IL-6 and TNF-α remained undifferentiable for all the groups studied at all time points tested p.i. (Fig. 5c).

The severity of the pathological changes associated with SARS-CoV-2 infection was much more prominent in the lungs of control infected animals than in other groups on both days 4 & 14 p.i. Lung tissues of control-infected animals had more alveolar and peribronchiolar infiltration and epithelial desquamation when compared to the lung tissues obtained from other groups. Animals that received IMT-CVAX + AddaVax had significantly reduced lung pathological conditions, such as cellular infiltration, compared to control-infected animals. On day 14 p.i., female hamsters injected with IMT-CVAX + AddaVax had a minor lung pathology than any other group studied (Fig. 5d). These data corroborate our clinical findings and establish the in-vivo efficacy of our candidate vaccine IMT-CVAX in restricting viral replication thereby conferring protection among the vaccinated animals.

### IMT-CVAX immunization in hamsters resulted in the elicitation of anti-spike IgG and anti-sera confers neutralization of live SARS-CoV-2

The primary objective of a vaccine intervention to combat viral infection is to elicit high titer of neutralizing-type antibodies. Indeed, the IMT-CVAX vaccinated groups elicited higher levels of anti-spike (RBD) specific IgG antibodies, which were also capable of neutralizing SARS-CoV-2 (Fig-5b, ii-iii). The live virus neutralization titer data showed that IMT-CVAX + AddaVax vaccinated groups have a decent NT50 value (1:969) and effectively neutralize the Hong Kong strain of SARS-CoV-2 with which challenge infection was carried out (Fig. 5d). In addition, hamster sera also neutralize Delta variant with similar extent (Fig. 5e). Taken together, these data provide clear evidence of the vaccine-induced correlates of protection that confers protection against SARS-CoV-2.

### IMT-CVAX immunization induces germinal center (GC) response in hACE2 transgenic mice

Long-lasting and high-affinity antibodies are produced in the microanatomical structures known as GCs found in secondary lymphoid organs. Extensive cross-talk prevails between Tfh cells and B cells during the multifaceted process of GC formation (35). Thus, a GC response indicates vaccine effectiveness in inducing optimal humoral immunity. To understand the formation of GCs in response to the spike-based vaccine, we measured the GC-B cells and Tfh cells after 2-dose immunization with IMT-CVAX adjuvanted with AddaVax. The vaccine induced a high titer antibody response after the second immunization dose (Adjuvant alone vs. vaccine + Adjuvant; p=0.0007). We observed that the adjuvanted vaccine was capable of inducing the GC B cells more significantly than the control group, both in frequency (Adjuvant alone; 0.53 ± 0.36, Vaccine + Adjuvant; 3.018 ±0.53, p=0.0159) and in count (Adjuvant alone; 471±222, vaccine + Adjuvant; 2133±619, p=0.0381). Because T cells play a crucial role in anti-viral response and in establishing humoral immunity, we also examined the ability of this vaccine to induce T cell responses. A significantly higher frequency of activated CD4+ T cells was found after 2-dose immunization both in frequency (Adjuvant alone; 2.61±0.29, Vaccine + Adjuvant; 3.88±0.29, p=0.0381) and in count (Adjuvant alone; 1158±159, Vaccine + Adjuvant 1796±101, p=0.0190). The specialized subset of CD4+ T cells, known as T_fh_ cells, is crucial for assisting B cells and establishing the GC response. Like GC B cells, a significantly higher frequency of CXCR5+PD1+ T_fh_ cells was induced in response to the vaccination (adjuvant; 0.1450±0.015, vaccine; 0.3833±0.10, p=0.0381). The results suggested that the IMT-CVAX could induce the T_fh_ cells and GC response which are the hallmarks of robust humoral immunity (Fig 6 d-k).

**Fig. 6:**
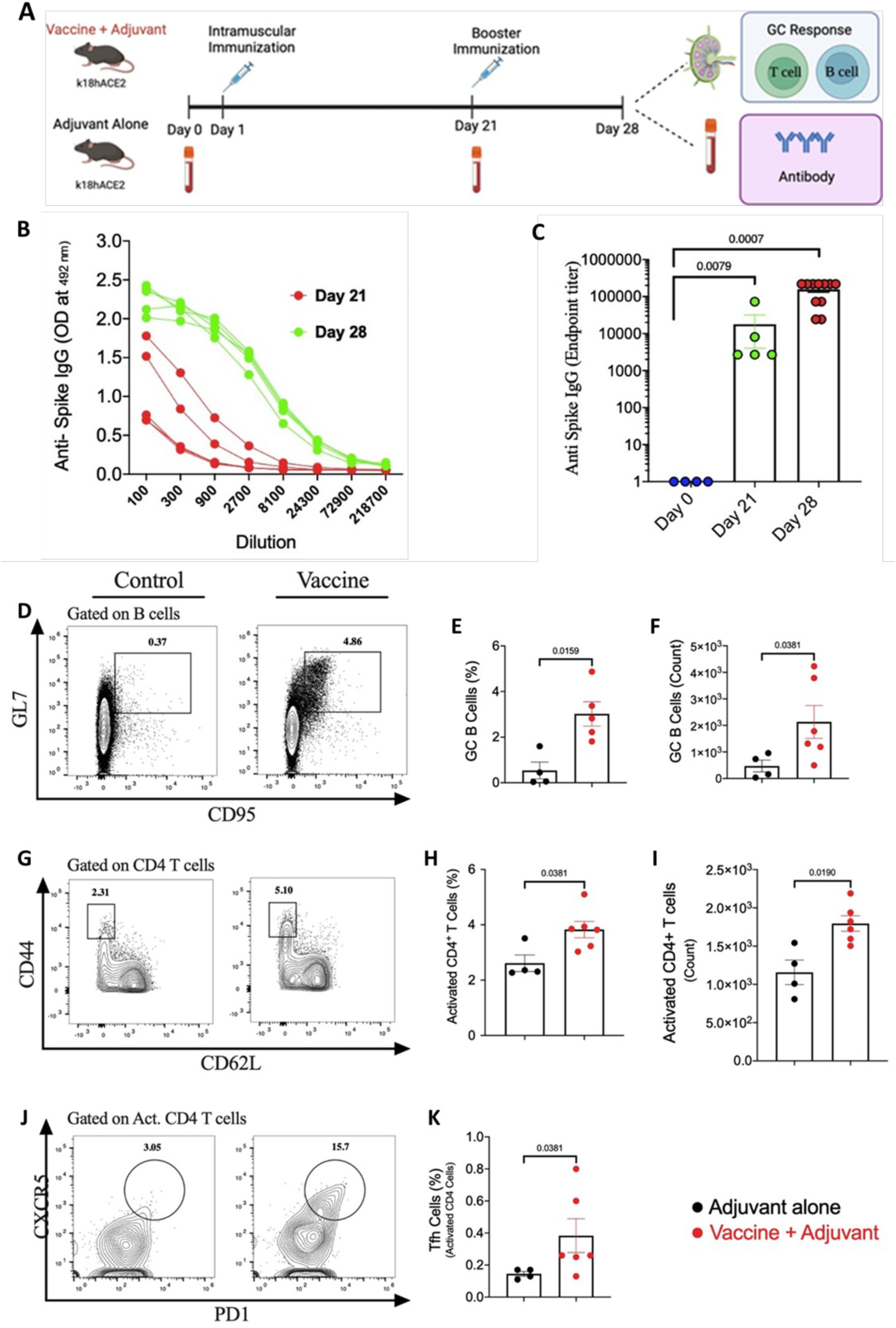
The 2-dose immunization with IMT-CVAX + AddaVax^TM^ was sufficient to induce germinal center (GC) response. (A) Study design to examine the ability of vaccine in inducing the GC response in hACE2-Tg mice. (B) Curves presenting the Spike-specific IgG in 3-fold serially diluted serum samples tested at pre- and post-booster immunization. (C) The end-point titre (ET) of SARS-CoV-2-Spike specific IgG at different time points post-immunization. (D) Representative plots depicting the detection of GC B cell subsets in total live B cells in vaccine and control group at day 28. Comparison of GC B cell in vaccine and control group for (E) frequency and (F) count. (G), Representative FACS plots showing the activated CD4+ T cells in vaccine and control group. Comparison of activated CD4+ T cell among the total live CD4+ T cells for (H) frequency and (I) count, in vaccine and control group. (J) Representative FACS plots depicting the detection of Tfh cells among activated CD4+ T cells in vaccine vs control group. (K) Comparison of Tfh cell frequency among total CD4+ T cells in vaccine vs control group. Black bars indicate mean ± s.e.m. Schematic in A was created using Biorender. Statistics by two-tailed Mann-Whitney test.

## Discussion

Equipped with the arsenal of evolving differently-infective and vaccine-immunity-evading variants, SARS-CoV-2 continues to cause significant morbidity and mortality worldwide (1). For maximum possible containment of SARS-CoV-2, the current situation warrants a continuous supply and rolling-out of vaccines with broad and durable protective efficacy. The recent COVID-19 pandemic has also taught global scientific communities and industries a lesson about the need for production-ready vaccine technology platforms for the swift response to potential future pandemic(s). Protein-subunit-based vaccines possess a long, proven history of safety and efficacy, and therefore we have adopted this path for developing a vaccine candidate against SARS-CoV-2. The relevance of prefusion stabilized spike protein-subunit as the choice of a vaccine targeting the SARS-CoV-2 surface glycoprotein has been highlighted in the early days of the COVID-19 pandemic ramification (17, 36, 37). However, the inherent instability, complex nature, and dynamic conformational architecture of spike protein is a significant challenge to be addressed before using it as a potential antigen for vaccine development (17). The discovery of protein-engineering strategies to stabilize the antigenic prefusion conformation of the SARS-CoV-2 spike by different amino acid substitutions in the native protein sequence was a significant milestone (17). These strategies include the introduction of disulfide bonds between nearby cystines, substitutions to fill the cavity by increasing the chain length of specific amino acids, inserting different numbers of proline amino acids (e.g., 2P, 6P-hexa-pro) in the loop region of the protein structure, etc. (17, 19, 36).

In our present endeavor, we developed a next-generation prefusion stabilized trimeric spike protein subunit-based vaccine candidate named IMT-CVAX to prevent COVID-19 infection. The excellent pre-clinical immunogenicity and protective efficacy conferred by IMT-CVAX strengthen the suitability of use of prefusion stabilized spike protein subunit-based vaccine for clinical development. The candidate antigen is designed to express prefusion stabilized trimeric spike ectodomain (ECD) protein in the CHO-S cell line. To stabilize the protein in its prefusion state, we have made six prolines (hexa-pro) substitutions in the native SARS-CoV-2 spike protein D614G sequence and modified the furin cleavage site, as described earlier (17).

In the other approved recombinant vaccines, two proline amino acid substitutions (2P) have been done for the prefusion stabilization of spike protein. In a recent study, the animal immunogenicity to a different version of the spike, i.e., native, 2-P, and 6-P, has been systematically compared. It was reported that using six proline substitutions (hexa-pro) in a spike increases its immunogenicity and efficacy and enhances the protein’s expression in recombinant systems. This study revealed that i) spike with 2-P substitution elicits a better immunogenic response when compared with native spike, ii) the 6-P substituted spike antigen was found to be more immunogenic than 2-P, and iii) a better T-cell response and cross-neutralization of VoCs was also observed upon immunization with 6-P spike, than 2-P (38). This data confirms that hexa-pro (6-P) pre-fusion stabilized spike could be a better and broadly-protective vaccine antigen. The choice of using the spike ECD as an immunogen in our study, rather than only RBD, was based on the fact that there are specific cryptic and conserved neutralizing epitopes in the non-RBD regions of the spike, which might be advantageous and could offer larger protective epitope-diversity, thereby expanding the vaccine’s range-of-protection to other VoCs that are already circulating and/or might arise in the years to come (39).

The CHO cell line has been used to produce recombinant biopharmaceuticals, including vaccines and biotherapeutics, since 1986 (40). One of the characteristic features of CHO cell lines is their ability to provide human-like post-translational modifications (PTM) in the molecule, which is particularly important in the context of spike protein as it is profusely glycosylated. In addition, CHO cell lines are capable of high cell-density growth and high productivity, making them one of the most favored expressions hosts for producing complex recombinant proteins at an industrial scale (41). We recently demonstrated the suitability of the CHO-S suspension cell line in combination with expression vector pCHO 1.0 for the high-yield expression complex protein like glycosylated monoclonal antibodies (20). However, this is for the first time that we are reporting the applicability of this cell line-vector combination for the expression of prefusion stabilized trimeric spike protein (IMT-CVAX) and its successful demonstration as a viable vaccine candidate against SARS-CoV-2 using pre-clinical models. Using an innovative bioprocess approach, we optimized the protein expression process and achieved very high yields of trimeric IMT-CVAX using bioreactor-based scale-up. It is expected that the expression of IMT-CVAX protein could be further manifold increased in large-scale bioreactor based scale-up in an industrial setup.

In some previous studies, different versions of CHO cell lines have been used to express native and prefusion-stabilized spike protein. Schaub et al. has demonstrated the feasibility of the expression of trimeric His8-TwinStrep-tagged hexa-pro (6P) version of spike using the ExpiCHO cell line and achieved >30 mg/L of protein expression (42). In another study, the CHO-S cell line was used to express his-tagged trimeric spike protein with 2P substitutions (K986P and V987P). It achieved a maximum expression yield of 53 mg/L using stable cell pools in fed-batch operation (43). Furthermore, Pino et al. have reported CHO cell-based expression and bioreactor based scale-up of the 2P variant of his-tagged trimeric spike protein and demonstrated the industrial feasibility of the process for producing high-quality spike antigen for diagnostic and potential for the vaccine (44). Noteworthy, during our study, we achieved very high expression of prefusion stabilized spike, close to 430 mg/l. To the best of our knowledge, this is the first time we are reporting a developmental process capable of generating biopharmaceutical grade hexa-pro (6P) prefusion stabilized trimeric spike protein ECD produced in cGMP banked suspension CHO cell line. However, most of the previous studies report the expression and purification of spike protein with one or other kind of affinity tags. This is to highlight that, the presence of affinity tags (e.g., His-tag) in final vaccine formulation could be a matter of concern from a regulatory compliance perspective. There are limited very studies on the purification of tag-free spike protein. In this context, we have developed a simple, scalable and robust purification process for the first time that uses non-affinity-based methods to obtain highly pure tag-free IMT-CVAX. This finding could fill the scientific and technological understanding gap for the purification of complex proteins like spike. Considering the high protein yield, the present study’s finding may also help develop affordable vaccines, particularly for low & middle-income and resource-poor countries.

The viability of mammalian cell culture during production is a critical parameter that determines the complexity of the overall protein-purification process. We observed that the purity of IMT-CVAX at the time of harvest was directly proportional to CHO-S cells’ viability during the production phase. Cultures having ≥85% VCD were easily purified using TFF and only one-step chromatography (either anion exchange or size exclusion). In cases where the VCD had dropped below 85%, two-step chromatography using anion exchange followed by sized-exclusion chromatography is required to obtain highly pure IMT-CVAX. Cell cultures with viabilities <85% contained more cellular protein impurities, which required an additional chromatography step for their complete separation. Prior to chromatographies, tangential Flow Filtration (TFF) played a pivotal role in IMT-CVAX purification as it separated the smaller cell-derived impurities and, therefore, highly recommended based on our current finding.

We performed detailed analytical analyses to acertain the structural and functional properties of IMT-CVAX. In reducing SDS-PAGE and western blotting analyses, IMT-CVAX appeared to have a single band of ∼180 kDa size, and no degradation in protein was observed, confirming the prefusion stabilization of the protein. The observed size of the monomer was in agreement with the predicted theoretical molecular weight of the spike monomer. This data was further confirmed by subsequent LC/MS analyses of the reduced IMT-CVAX. The trimeric nature of IMT-CVAX, corresponding to ∼ 660 kDa, was demonstrated by the SE-HPLC analyses done under non-reducing conditions and was confirmed by further DLS, AUC, DSF, and nsTEM analyses. The DSF results also hinted towards the potentially high thermostability of IMT-CVAX as it appeared to have a melting temperature of 64℃ (17). However, this is currently just a conjecture and needs further in-depth investigation. The ELISA data suggests that IMT-CVAX could have a strong affinity with the human ACE2 receptor (14). The overall analytical characterization results revealed that IMT-CVAX is in a homogenous, trimeric, prefusion-stabilized form and retains its conformational integrity.

Immunization of BALB/c mice and Golden Syrian hamsters with the AddaVax^TM^ adjuvanted IMT-CVAX induced a robust antibody response characterized by its broad spectrum of neutralization of the Wild Type, Alpha, Beta as well as Delta variant of SARS-CoV-2. Furthermore, the AddaVax™ adjuvanted vaccine also enhanced the induction of B-cells and CD4+ T cells in C57BL/6J and K18-hACE2 mice. The development of T_fh_ cells and GC B cells are crucial for the induction and maintenance of a high-affinity antibody response (32).

Based on the data collected from daily monitoring of body weight, clinical signs, and viral load and histopathological analyses, it was clear that the hamsters immunized with the AddaVax™ adjuvanted IMT-CVAX formulation were better shielded from SARS-CoV-2 associated clinical signs than the control-infected group. AddaVax™ is a squalene-based nano-emulsion of oil-in-water and has a composition similar to that of MF59 adjuvant. MF59 is a well-known adjuvant used in a licensed seasonal influenza vaccine (Fluad^®^) for more than 2 decades and is also part of the H1N1 vaccine (Focetria® and Celtura^®^). MF59 adjuvanted vaccines are well tolerated, safe, and effective in the elderly and young children (26). As previously reported, MF59 adjuvant acts via “depot effect,” which consequentially allows dose sparing of the antigen. It creates an immunostimulatory locale at the injection site, permitting slow antigen release leading to a durable immune response. It enhances the uptake of antigens by dendritic cells, leading to better seroconversion and cross-protection (45, 46, 26, 27). This could be why we observed a broad neutralization potential of IMT-CVAX immune sera against different VoCs of SARS CoV-2 during our study. The enhanced breadth of response is considered a valuable feature of SARS-CoV-2 vaccines to tackle emerging VoCs.

The strains of SARS-CoV-2 reported during the first wave of the pandemic were a major concern and targeted by most first-generation COVID-19 vaccines; however, several other VoCs have evolved during the subsequent waves of the pandemic. These VoCs exert a more serious public health threat, as they have evolved against vaccine selection pressures and therefore possess efficient capabilities of evading the human immune response (47). One such example is the Delta variant of SARS-CoV-2, which was the primary cause of COVID-related mortalities, after the Alpha variant, especially in the Indian subcontinent and other parts of the world. Studies have shown that this variant has a higher transmission rate and replicates faster than the other variants (48–51). More importantly, the Delta variant suppresses the host’s ability to generate an immune response, enabling it to thrive within the host for extended periods. Therefore, vaccines offering a more comprehensive range of protection against multiple VoCs, including Delta are the need-of-the-hour (47). Noteworthy, the neutralization assays we performed using the sera of immunized mice and hamsters indicated that antibodies generated against IMT-CVAX effectively neutralized not only the Wild-Type, but also cross-neutralized the Alpha, Beta and, more importantly, the Delta variant of SARS-Cov-2. This cross-neutralization gives us hope that IMT-CVAX may provide effective protection against any other variant(s) that may arise in the future. A prominent GC-response is the basis of this good quality and broadly protective humoral immunity. Our data on GC-response in K18-hACE2 mice further supports that the vaccine is capable of generating an optimal humoral immune response by inducing potent GCs. We have previously reported the signature of T-cell-based immunity of inactivated viral vaccine and established the assays (32).

In conclusion, our present research describes the development of a highly immunogenic and efficacious recombinant spike protein subunit-based adjuvanted vaccine candidate IMT-CVAX expressed in CHO-S cell line. The key findings of the present study are: 1) development of a high yield, robust, scalable and industry-compatible cell line and bioprocess for expression of prefusion-stabilized trimeric spike protein-based vaccine candidate IMT-CVAX (encoded by IMT-C20 gene), 2) optimization of very high level expression of spike protein in suspension-based mammalian cell line (CHO-S), 3) development of a robust process for purification of biopharmaceutical grade tag-free spike protein that can be used for research, pre-clinical, regulatory and clinical development, 4) excellent immunogenicity and pre-clinical efficacy of adjuvanted IMT-CVAX against SARS-CoV-2 challenge in hamster model, 5) use of MF59-like adjuvant that is capable of inducing both humoral and cellular immune response, desirable for broadly efficacious COVID-19 vaccines, 6) the anti-sera generated against the IMT-CVAX conferred excellent breadth of cross-protection against different VoCs of SARS-CoV-2 with similar magnitude, and 7) IMT-CVAX was found to induce robust Tfh cells and germinal centre responses that are pivotal for the generation of a long-lasting vaccine-specific immune response and, 8) the present expression platforms can be used for development of next generation affordable vaccines against SARS-CoV-2 and, could also be helpful in developing vaccine candidates against other related pathogens.

## Material and Methods

IMT-C20 was designed using the native Spike sequence. It was synthesized in pcDNA 3.4, and its expression was checked by transient production in ExpiCHO cells. IMT-C20 was then cloned in Freedom pCHO 1.0 and CHO-S cells were transfected with this construct. In the presence of dual selection pressure, four IMT-CVAX producing stable CHO-S cell lines were generated, and out the four, the cell line with highest expression (C20-2a) was selected for further optimizations of feed and temperature conditions during production phase at flask level. Bioreactor level scale-up was performed using the flask-level optimized conditions. The purification process of IMT-CVAX was optimized, and subsequently, pure IMT-CVAX was characterized using various biochemical and biophysical techniques. Mice and hamster models were used to assess the immunogenicity and pre-clinical efficacy of IMT-CVAX, respectively. Details of all the procedures as mentioned above are elaborated in *SI Appendix*.

## Acknowledgments

This study is supported by funding from the Council of Scientific and Industrial Research (CSIR), New Delhi, India, under project code MLP054. ST acknowledges funding support from DBT-BIRAC (BT/CS0007/CS/02/20) and Crypto Relief Funds (SP/CRRE-21-0004.05) for the Hamster challenge study. The work carried out at the National Institute of Immunology (NII) was supported by the Biotechnology Industry Research Assistance Council, Department of Biotechnology, India grant (BT/COVID0010/01/20) to NG. We thank Neeraj Khatri from IMTECH Center for Animal Resources and Experimentation (iCARE) for his support in facilitating the supply of mice for the immunogenicity experiment. We thank Inderjit Singh and Sudipta Das for their help in maintaining the K18 hACE2 line at the small animal facility of the NII. We thank Anil Kaul (Member, CSIR-Scientific Advisory Board), Deepak Sharma, Ashish and Rajeev K. Tyagi from CSIR-IMTECH for providing scientific and technical inputs in the project. We thank the CSIR-COVID Strategic Group (CSG), New Delhi: Ram Vishwakarma, Sanjeev Khosla, Rakesh Mishra for review of the program and scientific inputs, and administrative support during the execution of this project. Administrative supports from R.P. Singh and Nidhi, CSIR Innovation Management Directorate is acknowledged.

## Author Contributions

RPNM conceived and supervised the overall study wrote and reviewed the paper; Jitender, BVK, Sneha S, and RK performed cloning, expression, scale-up, purification, and characterization of antigen and wrote the manuscript draft; GV, PMM, Pooja performed mice immunogenicity and ELISA; Mumtaz, NK, DSM, AKB, and Shubham S performed protein purification and analytical assays; SK and RPR performed pseudovirus neutralization assays using mice sera; SKN, CN, CMJ, and ST performed hamster studies; BN, DS, and NG performed hACE2-Mice and T-cell Immunology.

## SI Appendix

### Materials and Methods

#### 1. Cell Lines, Media, and Vectors

cGMP banked CHO-S™ cell line, Freedom pCHO 1.0™ mammalian expression vector, lipofectamine™ transfection reagent, transfection enhancer, CD FortiCHO™ medium, Glutamine, ActiCHO™ P medium, ActiCHO™ feed A & feed B were procured from Thermo Fisher Scientific, USA. All the media and feeds used in the study were chemically defined and protein/serum-free to ensure compatibility with industrial quality standards set for the process and product.

#### 2. Design of Antigen and Synthesis of Gene Construct

The gene encoding the native SARS-CoV-2 spike protein (GenBank Accession Number: MN908947) was used as a template to design the IMT-C20 construct (used for expression of recombinant SARS-CoV-2 spike protein, IMT-CVAX). The native signal sequence at the N-terminus (1-13 amino acid) was replaced with a 23 amino acid long heterologous secretory signal peptide (murine immunoglobulin κ-chain) and inserted D614G substitution. Furthermore, removal of transmembrane domain and cytoplasmic tail at the C-terminus ( from 1196-1273 amino acids), and insertion of T4 fibritin foldon trimerization motif and 8x-histidine at C-terminus was done. Furin cleavage site’s amino acids RRAR were substituted with GSAS residues and 6 Proline substitutions (F817P, A892P, A899P, A942P, K986P,V987P) were made as described by Hsieh et al (10). The IMT-C20 was placed between the CMV promoter and WPRE site of pcDNA 3.4 (Fig-1d) and custom synthesized with the help of GenArt, Thermo Fisher, USA.

#### 3. Transient Expression and Purification of His-tagged IMT-CVAX

##### a) Expression of His-tagged IMT-CVAX

ExpiCHO™ cells were transfected with pcDNA 3.4–IMT-C20 (final concentration of ∼0.8 µg/mL) using the Lipofection kit (Invitrogen, USA), which consisted of ExpiFectamine™ CHO reagent and OptiPRO™ SFM (Serum Free Media) and were added as per the manufacturer’s instructions. For one transfection reaction, with 50 mL working volume, 40 µL plasmid DNA (1 mg/mL stock) was diluted with 2 mL OptiPRO™ SFM in a 50ml Falcon tube. In another 50 mL falcon tube, 160 µL ExpiFectamine™ CHO reagent was diluted with 1.84 mL OptiPRO™ SFM and incubated at room temperature for 5 minutes. Subsequently, the diluted ExpiFectamine™ CHO reagent was added to the diluted plasmid DNA and incubated for 5 minutes at room temperature. Slowly and in a dropwise manner, the ExpiFectamine™ CHO reagent and plasmid DNA mixture was added into a shake flask containing cells with continuous and gentle swirling.

The transfected cells were incubated in a CO_2_ shaker incubator at 32℃, 5% CO_2_, 125 rpm and ≥80% relative humidity for 24 hours. Approximately 18-22 hours post-transfection (i.e., Day 1), 300 µL ExpiFectamine CHO Enhancer and 8 mL ExpiCHO Feed (Max Titer, Thermo Scientific, USA) were added the culture and incubation was continued. Daily 1mL samples were collected for analyses. Five days post-transfection, 8 mL ExpiCHO™ Feed (Max Titer) was added, and the culture was further incubated. The samples were used to determine the transfected cells’ viability and viable cell density (VCD). The culture was harvested when the viability dropped below 80% (typically between 12 to 14 days post-transfection). The cells were separated from the media by centrifugation at 4000 rpm at 4℃ for 20 min. Protein expression in the daily collected samples was checked by reducing SDS-PAGE and western blotting.

##### b) Purification of His-tagged IMT-CVAX

The culture was harvested, and the broth was collected by centrifugation. TFF was performed using a 30KDa MWCO (Molecular Weight Cut-Off) membrane (Pall Corporation, USA). The spent media was first concentrated via TFF, and then the components of the spent media were exchanged with the binding buffer (50mM Sodium Phosphate, 300mM NaCl, 10mM Imidazole, pH 7.4). At least 5 diafiltration volumes were exchanged with the chromatography buffer. The retentate from TFF was recovered and loaded onto the chromatography column containing 10 mL affinity resin (TALON Superflow containing Cobalt, Takara Bio, USA). A thorough wash followed the binding step to remove any undesired and unbound impurities, and then the bound proteins were eluted using elution buffer (50mM Sodium Phosphate, 300mM NaCl, 150mM Imidazole, pH 7.4). Elution was initially done under a gradient of 0-50% in 30 min and then directly at 100% elution buffer. All the collected fractions along with the binding pass-through, were analyzed by reducing SDS-PAGE (10% polyacrylamide gel).

#### 4. Development of CHO-S cell line for stable expression of IMT-CVAX

##### a) Sub-cloning of IMT-C20 into Freedom pCHO 1.0 Mammalian expression Vector without the 8x-His

IMT-C20 was PCR-amplified from pcDNA3.4 using the gene-specific primer pair C20-AvrII-Fw (Forward: CCTAGGGCCACCATGGAAACCGATAC) and C20-PacI-Rev (Reverse: CTTAATTAACTACTATTGCTCGTACTTCCCC). The restriction enzymes AvrII and PacI were incorporated into the forward and reverse primers, respectively, so that the gene can be cloned into Freedom pCHO 1.0. Phusion™ High-Fidelity DNA Polymerase (New England BioLabs, USA) was used for amplification. The amplified PCR product was run in agarose gel (1%). Approximately 2.9 kb band corresponding to the untagged IMT-C20 was excised from the gel and purified using the Monarch DNA Gel Extraction Kit (New England Biolab-NEB, USA). The gel-purified IMT-C20 and the empty Freedom pCHO 1.0 vector were double digested using AvrII and PacI (NEB, USA) restriction enzymes following the manufacturer’s instructions and were resolved on 1% agarose gel. The gene and vector bands were excised from the gel and purified using a DNA gel extraction kit (NEB, USA). Subsequently, the tag-free IMT-C20 was ligated into Freedom pCHO 1.0 vector using the Quick Ligation Kit (NEB, USA), following the instructions provided.

##### b) Transformation of *E. coli* DH5α using the ligated Freedom pCHO 1.0–IMT-C20 Construct

The chemically competent *E. coli* DH5α cells (Invitrogen, USA) were transformed with the ligation product (Freedom pCHO 1.0+IMT-C20) and spread on Luria-Bertani (LB) agar plates containing Kanamycin (50µg/mL). The plates were incubated at 37℃ for 12-16 hours, after which a single colony was picked up and inoculated into 10 mL LB broth containing Kanamycin (50µg/mL). The culture was incubated at 37℃, 150 rpm, and grown till the mid-log phase (OD _600_= 1). Subsequently, 1 mL of culture was re-inoculated into 200 mL LB broth with Kanamycin (50µg/mL). This culture was incubated overnight at 37 ℃, 150 rpm. The culture was harvested after 12-16 hours and was centrifuged at 8000 rpm, 4℃ for 10 minutes to separate the cells from the broth. The plasmid (Freedom pCHO 1.0–IMT-C20 was purified from the cell pellet using EndoFree Plasmid MaxiPrep™ Kit (Qiagen, USA) following the manufacturer’s instructions. The concentration of the isolated plasmid was determined using a UV-VIS spectrophotometer (NanoQuant Infinity M200 Pro, Tecan Trading AG, Switzerland) at 260 nm wavelength. To confirm that IMT-C20 ligation in Freedom pCHO 1.0, the construct was digested using AvrII and PacI followed by agarose gel electrophoresis (0.8% agarose gel).

##### c) Transfection of CHO-S cells with Freedom pCHO 1.0–IMT-C20 Construct

The CHO-S cells were grown in Complete CD FortiCHO medium and sub-cultured in their mid-log phase (∼ 20 × 10^5^ cells/mL VCD) for 4 successive passages. One day before transfection, cells were passaged at ∼5 × 10^5^ cells/mL in Complete CD FortiCHO medium and incubated in a CO_2_ shaker incubator set at 130–150 rpm, 37°C, and 8% CO_2_. Before transfection, the cells were counted and diluted to get a VCD of ∼10 × 10^5^ cells/mL in a new flask with a 30 mL working volume. The Freedom pCHO1.0_MT-C20 construct was linearized using NruI restriction enzyme (NEB, USA), and transfection was performed as per the previously established protocol (1).

##### d) Development of IMT-CVAX expressing stable CHO-S cell line

A two-phase selection scheme was used to generate four different stable cell pools. The transfected CHO-S cells were grown under increasing concentrations of Methotrexate (MTX) and Puromycin in a complete CD FortiCHO medium. Forty-eight hours post-transfection, cells from the shake flask were seeded into 2 T-150 flasks at a cell density of ∼5 × 10^5^ cells/mL in different selection mediums. Puromycin (10ug/ml and 20ug/ml) and MTX (100nM and 200nM) were added. The T-Flasks were incubated in a static CO_2_ incubator set at 37℃, 8% CO_2_ and 95% relative humidity. Cell viability was checked daily. The cells were sub-cultured when the viability dropped to less than 85% (usually after day 7). Sub-culturing continued until the cells maintained viability ≥85%, marking the end of selection phase 1. Cells from selection phase-1, at a viable cell density (VCD) of 10 × 10^5^ cells/mL were sub-cultured for selection phase-2 and the criteria same as selection phase-1 were followed, except increased amounts of selection agents (30µg/ml and 50µg/ml of Puromycin and 500nM and 1000nM of MTX) were used. By the end of selection phase 2, four different stable cell pools (IMT-C20 1a, 1b, 2a and 2b) were generated. All four stable cell pools were banked, stored in liquid nitrogen and analyzed for expression. The detailed protocol for stable cell line development is already reported in our previous study (1).

#### 5. Optimization of conditions for stable expression of IMT-CVAX in the shake-flask and bioreactor

##### a) Shake-flask Optimization for Expression of IMT-CVAX

Fed-batch cultures at shake flask level were performed in 125 mL Erlenmeyer flasks with a starting volume of 30mL/flask and a VCD of 3 × 10^5^ cells/mL. Complete CD FortiCHO™ media (supplemented with 8 mM glutamine) was used with a 1% anti-clumping agent. Various feeding strategies were employed to assess their effect on cell growth and protein expression (Table 1). The feeding regimes with the best results were then subjected to a temperature shift from 37℃ to 32℃ (Table 2). The samples were taken at regular intervals for VCD, glucose and lactate analyses, viability%, and SDS-PAGE analyses.

##### b) Production of IMT-CVAX in cell culture bioreactor

Scale-up studies were carried out using the BioFlo 320 bioreactor (Eppendorf, Germany) to optimize the fed-batch production of IMT-CVAX with a 2L glass water-jacketed vessel. The batch was started with a working volume of 1L. The inoculum was prepared by thawing and reviving 1 vial of C20-2a (the selected clone corresponding to IMT-CVAX) stable cell pool first in 30 ml media and then subculturing into 100 ml and then 200 mL media (10% V/V) for the bioreactor. CO_2_ and O_2_ were used to control the pH and DO of the culture, respectively, during the bioreactor operation. All hardware, inoculum bottles, feed bottles, antifoam bottles, tubing, etc., were checked, assembled and sterilized by autoclaving. DO and pH probes were calibrated and sterilized by autoclaving. The vessel was filled with 1L 1xPBS (Phosphate Buffer Saline) and checked for any leaks by pressure hold test prior to sterilization by autoclaving. Post autoclaving, PBS was drained and the vessel was refilled with 900 mL medium under the biosafety cabinet and kept in stirring mode for 24 hours to ensure media sterility. Once the sterility and pressure hold run was completed and confirmed, 100 mL inoculum (3×10^5^ cells/mL VCD) was added to the bioreactor. During the bioreactor operation, samples were collected daily and checked for VCD, viability, glucose and lactic acid concentration profile. Data collection was done using the BioCommand software (Eppendorf, Germany).

#### 6. Optimization of tagless IMT-CVX purification

##### a) Clarification of the Culture Harvest by Centrifugation and Filtration

The culture was harvested and the cells were first separated from the broth by centrifugation at 4000 rpm for 20 minutes at 4℃. The supernatant was separated from the cell mass pellet and centrifuged at 8000-10000 rpm for 20 min at 4℃ to separate the remnant debris. Subsequently, the supernatant was filtered through a sterile 0.22 µm membrane filter within the biosafety cabinet to remove the minute cell debris and possible contaminants and stored at 4℃ till further use.

##### b) Buffer Exchange by Diafiltration Using Tangential Flow Filtration

The clarified broth was first concentrated and then exchanged with chromatography binding buffer, mentioned in the subsequent section, using a TFF cassette with a 30 kDa membrane cut-off (PALL Corporation, USA). At least five diafiltration volumes (DV) of the broth were exchanged before proceeding with chromatography. The retentate, containing the target protein, was collected and used in subsequent chromatography to purify IMT-CVAX.

##### c) Purification of IMT-CVAX by Ion-Exchange Chromatography

Anion exchange chromatography was used as an initial capture step. Initially, ANX and CaptoQ resins (Cytiva, USA) were tested for the capture chromatography at different pH (7.5, 7.0, 6.5, 6.0) on an AKTA chromatography system (Cytiva, USA). The TFF retentate was loaded and allowed to bind onto the columns, pre-equilibrated with the binding buffer (20 mM Sodium Phosphate, pH 7.5-6.0). The binding pass-through was also collected for evaluation. The bound proteins were eluted using a 0-100% gradient of elution buffer (binding buffer with 2M NaCl), and the peaks were collected. All the collected fractions and the binding pass-through were checked for the presence and quantity of spike protein.

##### d) Purification of IMT-CVAX by Size-Exclusion Chromatography

Size exclusion chromatography (SEC) was optimized using Sephacryl S300 HR 16/60 and Superdex 200 pg (Cytiva, USA) resins as the solid phase and Tris Buffer (20 mM Tris, 200 mM NaCl, pH 8.0) as mobile phase. The TFF retentate/pooled anion-exchange fractions containing IMT-CVAX protein were concentrated to ≤5mL using a 30 kDa MWCO centrifugal ultrafiltration device and loaded onto the columns. The individual fractions corresponding to different peaks were collected and analyzed by SDS-PAGE (10% polyacrylamide gel).

##### e) Determination of IMT-CVAX Concentration by densitometry

The protein content in samples from various batch and fed-batch cultures was done by gel-based densitometry using Bovine Serum Albumin (BSA, BioRad, USA) as a reference. The images of de-stained SDS-PAGE gels were obtained using a gel documentation system (Azure Biosystems Gel Imager C280 USA). The relative IMT-CVAX concentration in the samples was calculated against known concentrations of BSA with the help of UN-SCAN-IT gel V 6.1 software (Silk Scientific Corporation, USA).

##### f) Quantification of in-process Metabolites in the Samples Collected During the Bioreactor Run

Samples were collected daily to analyze metabolites (glucose and lactic acid) in batch and fed-batch cultures. Metabolite concentrations in the culture were determined using Ultra-Performance Liquid Chromatography (UPLC, Shimadzu Corporation, Japan) with a RID detector (RID-10A) and an Aminex HPX87-H column (BioRad, USA) in isocratic mode using 5mM H_2_SO_4_ as the mobile phase. The oven (CTO-10AS VP) was kept at 25°C with a pump flow rate of 0.6 mL/min. Before use, all buffers were filtered through a 0.22 µm filter and sonicated to remove any gas. All the samples were also filtered using 0.22 µm filters before injecting them into the column. The acquired chromatograms were compared to the standards, and glucose and lactic acid concentrations were determined using standard curves.

#### 7. Characterization of IMT-CVAX

##### a) Sodium Dodecyl Sulphate-Polyacrylamide Gel Electrophoresis (SDS-PAGE) and Western Blotting

SDS-PAGE was performed under reducing conditions, per the procedure reported elsewhere (2). Western blotting involved SDS-PAGE followed by transfer of the resolved protein bands from the gel onto nitrocellulose membrane (Sartorious, USA) using Trans-Blot® Turbo™ transfer system (BioRad, USA) at 2.0 A, 18-20V for 20 mins using transfer buffer (25 mM Tris, 190 mM glycine, and 20% methanol). Unsaturated sites on the blot were blocked by incubating it overnight at 4℃ in 5% skimmed milk. The blocked blot was washed thrice with 1X PBST before probing it with primary antibody, i.e., 1;1000 dilution of Anti-spike polyclonal antibody (SARS-CoV/SARS-CoV-2 Spike Protein Polyclonal antibody, Invitrogen, USA) in 1X PBST at room temperature for 3 hours with constant shaking. The blot was washed with 1X PBST 5 times and subsequently probed with Goat anti-rabbit IgG-HRP conjugate (1:20000 dilution in 1X PBST) at room temperature for 1 hour with constant shaking. The blot was washed 6 times with 1X PBST before developing it with ECL reagent and image acquisition was made using the gel documentation system (Azure Biosystems Gel Imager C280 USA).

##### b) Size-Exclusion High-Pressure Liquid Chromatography (SE-HPLC)

The SE-HPLC was performed to determine the approximate size of the intact IMT-CVAX using an SRT SEC-500 (Sepax Technologies, USA) column coupled to an HPLC system (Shimadzu Corporation, Japan). Samples were diluted and loaded onto the column, and isocratic elution was performed with phosphate buffer (100 mM sodium phosphate, 50 mM NaCl, pH 7.2). Samples were run at a 1mL/min flow rate, and detection was performed in a UV-Vis detector (SPD-20A) at 25 °C. LC Solution program (Shimadzu Corporation, Japan) was used to integrate the peak and calculate the proportion of aggregates in our samples.

Protein standards (Gel Filtration Markers Kit for Protein Molecular weights 29,000-700,000 Da, MWGF 1000-1KT, Sigma-Aldrich) were also run on the same column, under the same conditions. They were taken as a reference for the determination of the relative molecular weight of IMT-CVAX, based on the retention time.

##### c) Liquid Chromatography/Mass Spectroscopy (LC/MS)

The liquid chromatography/mass spectroscopy (LC/MS) system (Agilent 6550 iFunnel QTOF, Agilent, USA) was used to determine the intact mass of IMT-CVAX. The purified IMT-CVAX was diluted in reduction buffer (25 mM Tris, 25 mM NaCl, pH 7.5) to achieve a final 1mg/mL concentration. A concentrated TCEP solution was added to the diluted protein to a final concentration of 5 mM. The sample was incubated for 30 mins at 37°C before adding formic acid to a final concentration of 1%. The final concentration of protein samples used for LC/MS analyses was 0.2 mg/mL. C8 (150 × 3.0 mm, 300 Å) column was used with Buffer A (1% Trifluoroacetic acid in water) and Buffer B (1% TFA acetonitrile) as mobile phases. The sample volume loaded onto the column was 2 µl, and a linear gradient of Buffer B from 10%-70% in 15 min, 70–90% in 15-18 min, and 90%-100% in 18-19 min was applied at a flow rate of 0.4 mL/min and total ion chromatogram (TIC) was recorded. Agilent MassHunter, qualitative analysis software, was used to analyze the MS spectra, and the maximum entropy algorithm was used for the deconvolution of spectra.

##### d) Differential Scanning Fluorimetry

IMT-CVAX, at a concentration of 10µM, was mixed with 1% Sypro™ Orange dye (Invitrogen, USA) and was dispensed in PCR plates (50µL/ well), in triplicates. Sample containing only buffer (20mM, 200mM NaCl, pH 8.0) and 1% Sypro™ Orange dye and protein samples without Sypro™ Orange were taken as a negative control. The thermal denaturation profile was measured using Real-Time PCR (CFX 96, BioRad, USA) between 20℃ to 95℃ with a ramp rate of 0.5℃ per min. ROX filter was used to measure the fluorescent intensity, and the acquired data was analyzed using BioRad CFX maestro software and the first derivative of the data was used to calculate the T_m_.

##### e) Analytical Ultra-Centrifugation

The Beckman Coulter XL-I AUC™ was used to perform sedimentation velocity analytical ultra-centrifugation (SV-AUC) with a rotor speed of 40000 rpm for 100 scans with a three-minute time interval between each scan. AUC cuvette cells were filled with protein samples (IMT-CVAX) diluted into tris buffer (20 mM Tris, 200 mM NaCl, pH 8.0). In another cuvette, only Tris buffer was run that served as control. The SV-AUC run was initiated when the vacuum and temperature reached their set limits. The acquired data were analyzed using the SEDFIT™ software, and the results were plotted with the help of GraphPad™ Prism 8 software.

##### f) Circular Dichroism

Far-UV CD spectrum (195–260 nm) of IMT-CVAX (0.15 mg/mL) was measured on a spectropolarimeter (JASCO J-815 Jasco, Japan) using 1mm path length quartz cuvette at 20°C. Buffer (20mM, 200mM NaCl, pH 8.0) was used for baseline correction and the accumulation of three spectra was recorded. The data were analyzed using BeStSel Webserver™.

##### g) Transmission Electron Microscopy

The negative stain-Electron Microscopy (nsEM) of IMT-CVAX was performed per the published protocol (3). The test samples were diluted to 100μg/mL in tris buffer containing 20 mM tris, 150 mM NaCl, pH: 8.0 with 10% glycerol and pre-incubated for 5 min. 5μl volume of sample was allowed to bind onto a glow-discharged, carbon-coated grid for about 10 min, followed by blotting, negative staining with phosphotungstic acid (PTA) and air drying. Images were obtained using an electron microscope (JEOL JEM-2100 Electron Microscope).

##### h) Dynamic Light Scattering

Zetasizer v7.11 (Malvern, UK) was used to measure the size of the IMT-CVAX protein. Samples were centrifuged at 10000 rpm, 4℃ for 20 mins before the run and loaded onto clear zeta cell cuvettes. The instrument was equilibrated with tris buffer at 25℃ for 2 minutes, and intensity was measured as a function of time. The instrument’s software determined z-average, particle diameter and PDI from cumulative analysis of the scattered intensity autocorrelation function.

#### 8. Assessment of IMT-CVAX – ACE2 Affinity by ELISA

The ELISA plate was coated with 100ul of ACE2 (diluted in 1xPBS) so each well had 1ug/ml (100ng) of ACE2. The plate was incubated overnight at 4℃. After 12-16 hours of ACE2 binding, the plate was removed and washed 3 times with PBST (1x PBS + 0.05% Tween-20). 200 µl of blocking solution (1% BSA in PBS) was added to each well, and the plate was incubated at room temp for 3 hours. The plate was rigorously washed 4 times with PBST (1x PBS + 0.05% Tween-20), and 100 µl of diluted His-tagged IMT-CVAX (concentration ranging from 1ug/ml with 12 serial dilutions in 1xPBS) was added to the wells, in triplicates. PBS was used as the negative control in triplicates. The plate was incubated at room temperature for 3 hours and washed thoroughly 5 times with PBST (1x PBS + 0.05% Tween-20). Anti-His secondary antibody (Sigma A7058), at a 1/5000 dilution in 1% BSA in PBST, was added to each well, and the plate was incubated at room temperature for 2 hours. After the secondary antibody probing, the plate was washed 6 times with PBST. 100 µl of HRP substrate, TMB (Sigma Cat No. T4444) was added to the plate and incubated for 15 minutes at room temperature in the dark. Absorbance was taken at 655 nm. 1N H_2_SO_4_ was added over the TMB to stop the reaction, and immediately after, absorbance at 450nm was taken. The data obtained were plotted using GraphPad Prism 8.

#### 9. Assessment of Immunogenicity of IMT-CVAX in Mice Model

Immunogenicity study in mice model was conducted at IMTech Centre for Animal Resource and Experimentation (iCARE), CSIR-IMTech, Chandigarh, India. BALB/c mice (4-6 weeks old; mixed gender; n=8) were used for immunogenicity assessment of IMT-CVAX as per the protocol approved by the Institutional Bio-Safety Committee (IBSC) and Institutional Animal Ethics Committee (IAEC). IMT-CVAX was formulated with three different adjuvants: AddaVax, (a squalene-based oil-in-water nano-emulsion, similar to MF59 adjuvant), Alum (2% Alhydrogel) and Alum+CpG ODN 2006 (synthetic immunostimulatory oligonucleotide-ODN containing unmethylated CpG dinucleotides. Formulations were prepared according to the previously published procedures (4, 5). A 50μl vaccine formulation containing 10µg IMT-CVAX and respective adjuvants was injected via intramuscular route per mouse on days 0 (dose-1) and 28 (dose-2). The control group of mice received only formulation buffer (20 mM tris, 150 mM NaCl, pH: 8.0). Blood was collected from anesthetized mice on day 28 via retro-orbital plexus bleeding (post-dose 1 sera). On day 38 (10 days after dose-2 immunization), blood (post-dose-2) was collected from anaesthetized animals via cardiac puncture.

##### a) Ethics Statement

Experimental mice were housed in individually ventilated cages (IVCs) under controlled conditions of temperature (24–25°C), light (photoperiod of 12:12) and relative humidity (30–70%). Throughout the experiment, the animals were provided with water and synthetic pelleted feed *ad libitum*. The experimental protocols were approved by the IAEC of CSIR-IMTECH (Approval number IAEC/20/22), National Institute of Immunology (Approval number IAEC 556#20) and were carried out in compliance with the guidelines of the Committee for Supervision of Experiments on Animals (CPCSEA), Ministry of Fisheries, Animal Husbandry and Dairying, India.

##### b) Titration of Anti-Spike Protein Binding Antibodies by ELISA

Sera from mice were separated from clotted blood samples by centrifugation before titrating them by indirect ELISA. High-binding 96 well ELISA plates (ThermoFisher Scientific) were coated with 1 µg /mL (100 ng / 100µL / well) purified spike protein diluted in PBS and incubated at 4℃ overnight. The coated plates were washed three times with 1X PBST washing buffer (PBST: 1 × PBS + 0.05% Tween-20) using an ELISA plate washer (ELx50, BioTek, USA). Unsaturated sites of the coated ELISA plate were blocked by dispensing 200µL blocking solution containing 1% BSA in PBS and incubating at room temperature for 3 hours. The blocked ELISA plates were washed four times with 1x PBST. Subsequently, mice sera from vaccinated and control groups were subjected to 9 serial, two-fold dilutions from 1:100 to 1:25600 such that the ELISA plates received 100µL/well diluted mice sera. Primary antibody probing was done using sera obtained from immunized mice (having anti-IMT-CVAX IgG), at room temperature for 2 hours. The ELISA plates were then washed 5 times with 1X PBST. At room temperature, Goat anti-mouse IgG–HRP conjugate (1:20000 dilution; 100µL/well) (Sigma, USA) was used as the secondary antibody for 1 hr. After the secondary antibody probing, 100µL TMB substrate (Sigma, USA) was added to each well and incubated for 15 minutes, at room temperature, in the dark. The reaction was stopped by adding 100µL stop solution (1N H_2_SO_4_) to each well. Absorbance was measured at 450 nm using an ELISA plate reader (BioTek, USA). The ELISA plate readouts were plotted and analyzed using GraphPad prism™ software.

##### c) Titration of SARS-CoV-2 Specific Antibodies by Pseudovirus Neutralization Assay

To evaluate the *in-vitro* potency of immune mice sera, a SARS-CoV-2 pseudovirus neutralization assay was performed as per the protocol described in our previous publication (6). Pseudoviruses were prepared by transiently transfecting HEK293T cells with SARS-CoV-2 spike protein-expressing plasmid vector and a molecular clone pHIV-1NL4.3Δenv-nanoLuc which expresses all the structural proteins of HIV except Env-glycoprotein and a reporter nanoLuc luciferase enzyme (Promega, USA). Co-transfection of these two plasmid DNA vectors leads to the generation of pseudovirus particles decorated with SARS-CoV-2 spike protein on the surface and capable of a single entry into the cell. The neutralization assays were set up in 96-well plate format in which mice sera were serially diluted in 50µL growth medium, mixed with 50µL pseudovirus and incubated for 1 hour at 37°C. After incubation, 1×10^4^ cells (293T-hACE2-TMPRSS2) in 100µL was added and the assay plate was incubated at 37°C for 48 hours. Infection of the indicator cell line with pseudovirus was measured in terms of relative luminescence unit (RLU), and percent neutralization was determined with reference to the control in which no serum was added to the pseudovirus. Neutralization titres (Median infective dose; ID_50_) were determined as the serum dilution at which infectivity was reduced by 50% by using GraphPad Prism™ software.

#### 10. Assessment of Immunogenicity and protective Efficacy of IMT-CVAX in Hamster Models

##### a) Animals

Golden Syrian hamsters (mixed gender; 5-6 weeks old) (obtained from Biogen Laboratory Animal Facility, Bengaluru, India) were used for assessing the immunogenicity and protective efficacy of the IMT-CVAX vaccine using on vaccination–challenge approach. The animals were housed as separate groups (n = 14) and maintained in IVCs at 23±1°C temperature and 50±5% relative humidity, given access to standard pellet feed and water *ad libitum* and maintained on a 12-hr day/night light cycle at the viral biosafety level-3 facility, Indian Institute of Science, Bengaluru, India.

##### b) Immunization of Golden Syrian Hamsters

IMT-CVAX vaccine was formulated with AddaVax™ adjuvant, at a ratio of 1:1, just before use. The formulated vaccine had 100 µg spike protein / mL. One day before vaccination, all the animals recruited for the experiment were weighed and bled retro-orbitally for pre-immune sera collection. The groups of animals (n=14/group) were immunized with 100µL of adjuvanted vaccine formulations containing 10µg of IMT-CVAX intramuscularly (in the right hind limb region). Three doses of IMT-CVAX were given 2 weeks apart. Uninfected and infected controls group of animals (n=12), were also kept in the study and served as control. Total animal body weight was recorded before the administration of each vaccine dose. Post-immune sera were collected one day before infection at the end of week 6 from the first vaccine dose. The study scheme is provided in Figure 5A. Male and female hamsters received two doses of vaccine formulation according to the different groups, each at 3-week intervals before the virus challenge. Sera were collected at different time points as indicated. Lungs were harvested on day 4 or 14 post-infection (p.i) for estimating viral RNA load and histopathology analysis.

##### c) Challenge of Immunized Hamsters with Virulent SARS-CoV-2

Virulent SARS-CoV-2 was propagated for hamster challenge (Isolate Hong Kong/VM20001061/2020, Cat No.: NR-52282, BEI Resources, NIAID, NIH) (at 0.01 MOI) and titrated by plaque assay in Vero-E6 cells. Hamsters under intraperitoneal Ketamine (150mg/kg, Bharat Parenterals Limited) and Xylazine (10mg/kg, Indian Immunologicals Ltd) anaesthesia were challenged via intranasal route with ∼1×10^5^ plaque-forming units (PFU) of SARS-CoV-2 in 100µL PBS. The challenged hamsters were observed daily for clinical signs and weighed to record their body weight. On day 4 p.i., 7 hamsters from IMT-CVAX vaccinated group and 6 animals from the control groups were euthanized using an overdose of Xylazine. The remaining hamsters have monitored daily up to day 14 p.i. for body weight and clinical signs assessment. On day 14 p.i., the remaining hamsters were euthanized, and post-challenge sera were collected and stored frozen at −20°C until analyzed. The lung samples from both lobes were harvested for virological (left lobe) and histopathological analyses (right lobe).

##### d) Examination of Clinical Signs

Following the virulent SARS-CoV-2 challenge, the hamsters were observed daily for the following clinical signs and scored based on severity: Lethargy (none=0, mild=1, severe=2), piloerection (none=0, mild=1, moderate=2, severe=3), abdominal respiration (none=0, mild=1, severe=<u>2</u>), hunched back (none=0, mild=1, severe=<u>2</u>). Loss of body weight was also considered a clinical sign, with scoring done from a scale of 1 to 3 based on the per cent decrease in the body weight of animals (1-5 %= 1; 5.1-10%= 2; 10.1-15%=3). The above-mentioned clinical signs were recorded up to day 14 p.i. for all the groups.

##### e) Scoring of Lung Histopathology and Histopathology

The lung tissue samples were weighed and scored for focal and diffused hyperaemia. Based on the severity of hyperaemia, scoring was given on a scale of 1 to 3 (mild=1, moderate=2, severe=3). Lung images of each hamster from every group were also taken. Lung specimens of hamsters were fixed in 4% paraformaldehyde in PBS and embedded in paraffin blocks. Tissue sections of 4-6 μm thickness were stained with Haematoxylin and Eosin (H&E). The stained lung tissue sections were examined by light microscopy for three histological criteria (Alveolar infiltration and exudation, vasculature inflammation and peribronchiolar infiltration with epithelial desquamation). Based on the severity, each criterion was scored on a scale of 1 to 3 (mild=1, moderate=2, severe=3).

##### f) Quantification of Viral Load in Lung Tissues by qRT-PCR

Lung tissue samples from hamsters were processed using a tissue homogenizer, and total RNA was isolated using TRIzol (15596018, Thermo Fisher) as per the manufacturer’s instructions. A 10µL reaction mixture with 100ng of total RNA per sample in a 384-well block, was used to quantify viral RNA using AgPath-ID™ One-Step RT-PCR kit (AM1005, Applied Biosystems). The following primers and probes targeting the SARS CoV-2 N-1 gene were used. Forward primer: 5’GACCCCAAAATCAGCGAAAT3’ and Reverse primer: 5’ TCTGGTTACTGCCAGTTGAATCTG3’, Probe: (6-FAM / BHQ-1) ACCCCGCATTACGTTTGGTGGACC. The Ct values were used to determine viral copy numbers by generating a standard curve using the SARS-CoV-2 genomic RNA standard.

##### g) Quantification of Cytokine mRNA level by qRT-PCR

With the extracted total RNA, host mRNA expression of IFN-γ, TNF-α and IL-6 were quantified using suitable primers with 18S RNA as an internal reference gene. Briefly, cDNA was prepared from 1µg total RNA using PrimeScriptTM RT reagent Kit with gDNA Eraser (TaKaRa, USA). qRT-PCR was performed using an AgPath-ID 1-step RT-qPCR kit using 2.5μL of cDNA, 2.5μL of primer mix and 5μL of SYBR™ Green PCR Master Mix (Applied Biosystems). The following primers for hamster IFN-γ, TNF-α and IL-6 genes were used. IFN-γ: Forward primer: 5’TGTTGCTCTGCCTCACTCAGG3’ and Reverse primer: 5’AAGACGAGGTCCCCTCCATTC3’. TNF-α: Forward primer: 5’TGAGCCATCGTGCCAATG3’ and Reverse primer: 5’AGCCCGTCTGCTGGTATC3’ IL-6: Forward primer: 5’AGGCCATCCTGATGGAGAAG3’ and Reverse primer: 5’ GGTATGCTAAGGCACAGCAC3’. After running the RT-qPCR, fold change in gene expression compared to 18S RNA was calculated using the delta-delta Ct method. The mean of the Log_2_ fold change in gene expression for each gene in each group of animals was calculated.

##### h) Hamster Anti-Spike IgG Quantification by ELISA

The pre-infection sera from Syrian hamsters were used to quantify the anti-spike IgG by indirect ELISA method (Krishgen Biosystems). Briefly, SARS-CoV-2 spike-RBD protein pre-coated 96 well microplates were added with 100µL of the sample (1:1000 dilution) or standard (0-720ng/mL) and incubated for 1h at 37°C. After washing the plates 4 times with wash buffer, 100µL of goat anti-hamster IgG-HRP conjugate was added to each well and incubated for 1h at 37°C. The plate was washed and added with 100µL of 3, 3’ 5, 5’-Tetramethylbenzidine substrate and incubated for 15 min in the dark. The colour development was stopped by adding 2N H_2_SO_4_ and absorbance was read at 450 nm. The concentration of hamster IgG was automatically determined using a cubic spline-curve fit. The sensitivity of the assay kit was 12ng/mL.

##### i) Live Virus Neutralization Titre (NT) Assay

Live virus neutralization efficiency of post-immune sera was performed by the plaque reduction neutralization titer (PRNT) method. Briefly, 12-well plates were seeded with 0.2 × 106 cells/mL of VERO E6 and incubated for 48h at 37°C. All the sera samples were serially diluted 2-fold (1:10 to 1:320) in DMEM with 2 % FBS and then mixed properly with an equal volume of Wuhan or Delta strain SARS-CoV-2 (103 PFU/mL) and incubated for 1h at 37°C. A 100µL of this mixture was added in duplicates and incubated at 37°C for 1 h with shaking every 10-15 min. After incubation, the sample was removed completely, overlayed with 1 mL of 0.6 % avicel prepared in DMEM and incubated at 37°C with 5 % CO2 for 48 h. After incubation, avicel was removed completely and cells were fixed by adding 1 mL of 4 % formaldehyde in PBS. Then, cells were stained with a 1 % crystal violet solution. In each sera dilution, the number of plaques formed was counted and converted into per cent reduction in plaques using the below formula. A non-linear regression method was used to determine the sera dilution at which 50 % virus neutralization (NT50) occurred.

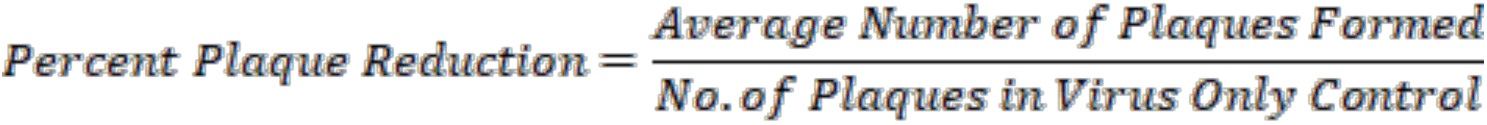

##### j) Statistical analyses

The ELISA and PRNT were analyzed using GraphPad Prism™ v 8.4.3 and represented as mean ± SEM. Statistical variations were determined by unpaired t-test (IgG ELISA and PRNT assay) or one-way ANOVA (Lung viral RNA load and cytokine mRNA level) or two-way ANOVA (Body weight, clinical signs and Lung histopathology) with Dunnett’s multiple comparisons tests. The values were considered significant when *P < 0.05, **P < 0.01, or ***P < 0.001.

#### 11. Humoral and Cellular Immune Response

##### a) Mice Immunization

The C57BL/6JK18-hACE2 mice were obtained from the Jackson Laboratories (USA) and maintained in a pathogen-free environment at Small Animal Facility, National Institute of Immunology, New Delhi, India. Five to six-weeks-old transgenic mice (C57BL/6J K18-hACE2) were immunized intramuscularly with 10 μg IMT-CVAX adjuvanted AddaVax™ adjuvant (InvivoGen) in 1:1 ratio. Blood samples and tissues were collected at different time points post-immunization, as illustrated in Figure 4 panel A.

**Fig. 4:**
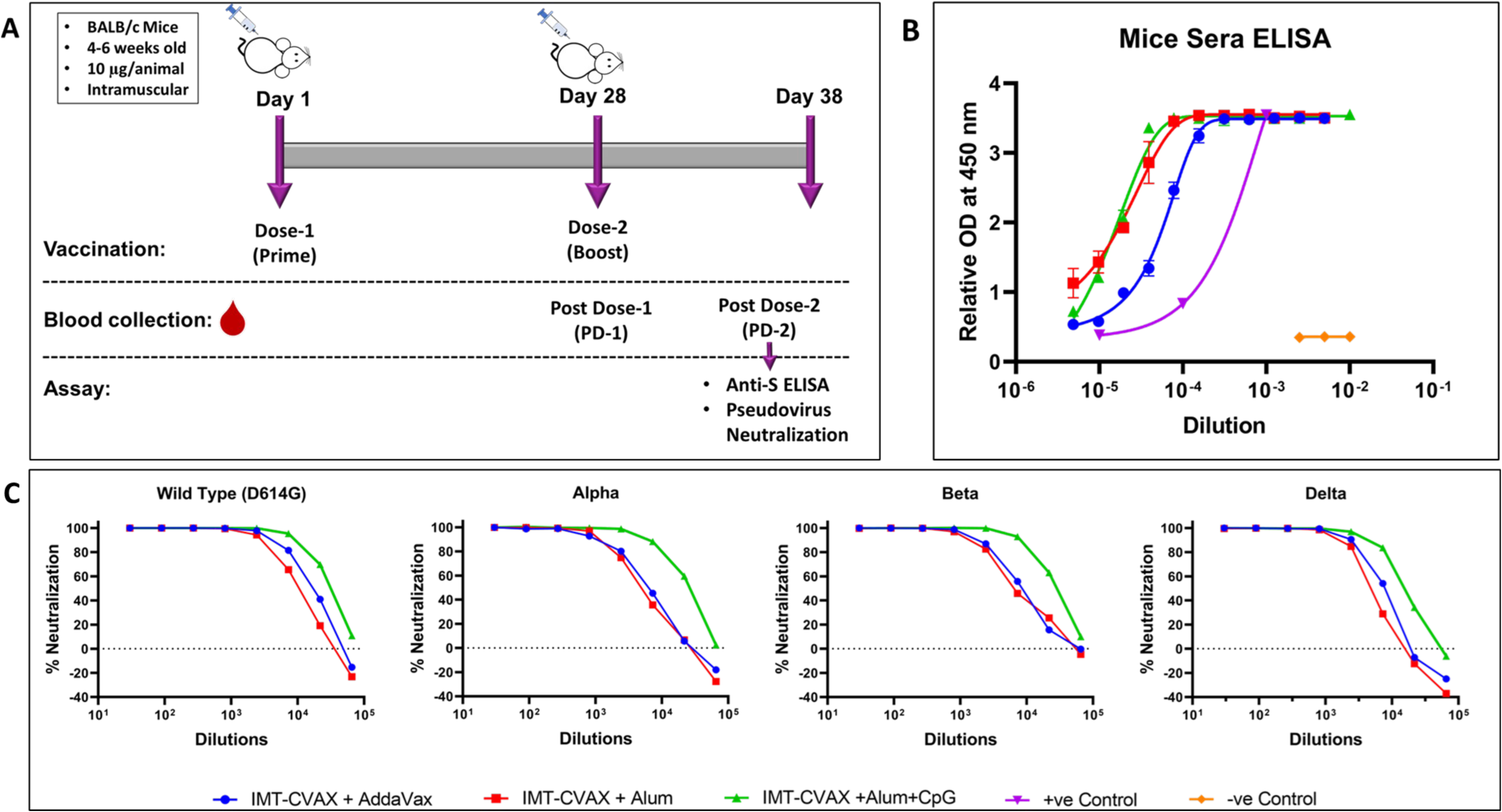
IMT-CVAX Immunogenicity in Mice. **(A)** is the process flow demonstrating the days of immunization, blood collection. **(B)** Serum samples showed positive results for the presence of anti-spike IgG. The legend is mentioned in Fig 4C. **(C)** Pseudovirus neutralization assays revealed that the sera obtained from immunized mice effectively neutralized the Wild-Type, Alpha, Beta and Delta SARS-CoV-2 pseudoviruses. Interestingly, a similar level of neutralization was obtained against various VoC, with slightly lower neutralization of Delta.

##### b) Flow-cytometry Analyses

Lymph node tissue samples were collected on day 28 post-booster vaccination for immunophenotyping. To detect subset-specific markers, the single lymphocyte suspension was stained with a panel of fluorochrome-coupled antibodies in FACS buffer (2 per cent FBS in PBS). The dead cells were excluded using the fixable viability dye eFluor 506 (eBioscience, Thermo Fisher, USA). Before staining with the respective panels, the cells were incubated with the Fc-block anti-CD16/CD32 (2.4G2) antibody. The GC B cells were measured after staining with anti-B220 PE-TR, anti-GL7 PE and anti-CD95 APC for 1 hr at 4°C. Activated CD4 T cells were stained using anti-CD4 AF700, anti-CD62L APC, and anti-CD44 FITC with the exclusion of anti-B220/45R PE-TR. The T_FH_ cells were measured using the surface markers anti-CXCR5 BV421 and anti-PD-1 PE by gating on the activated CD4 T cells. After incubation, the cells were washed and fixed with freshly prepared 1% paraformaldehyde (Sigma-Aldrich, USA). Data were acquired using BD LSRFortessa X-20 flow cytometer (BD Biosciences, USA) and subsequently analyzed using FlowJo 10.7.1, as previously described (Kalia A. et al., 2021).

##### c) ELISA

ELISA was performed as previously described (Kalia A. et al., 2021). Flat-bottom, 96-well ELISA plates were coated with 1 μg/mL SARS-CoV-2 spike protein in PBS followed by overnight incubation at 4°C. The coated ELISA plates were washed, blocked for 3 hrs at room temperature and again washed. Serum samples, serially diluted three-fold in blocking buffer (2% BSA and 0.05% Tween-20 in PBS), were added @ 50 μL/well and incubated for 2 hrs at room temperature. After incubation of the diluted sera, the ELISA plates were again washed and incubated for 1 hr with goat anti-mouse IgG – HRP conjugate (Southern Biotech, USA). The ELISA plates given a final wash were developed using o-phenylenediamine dihydrochloride peroxidase substrate (Sigma-Aldrich, USA). The optical density (OD) was measured at 492 nm wavelength using a MultiskanGO ELISA reader (Thermo-Fisher, USA). Sample diluent added into the Spike-antigen-coated wells was used as the blank. The mean of OD values obtained in the blank well in each plate was used for the background subtraction during analysis.

##### d) Statistical Analyses

Data were expressed as the mean ± SEM for all the experiments. The two-sided Mann-Whitney test was used to determine the significance of the differences between the groups. Version 9 of GraphPad Prism™ software was used to carry out statistical analyses. The ELISA and PRNT were analyzed using GraphPad Prism™ v 8.4.3 and represented as mean ± SEM. Statistical variations were determined by unpaired t-test (IgG ELISA and PRNT assay) or one-way ANOVA (Lung viral RNA load and cytokine mRNA level) or two-way ANOVA (Body weight, clinical signs and Lung histopathology) with Dunnett’s multiple comparisons tests. The values were considered significant when *P < 0.05, **P < 0.01, or ***P < 0.001.

---

**SI Fig. 1:**
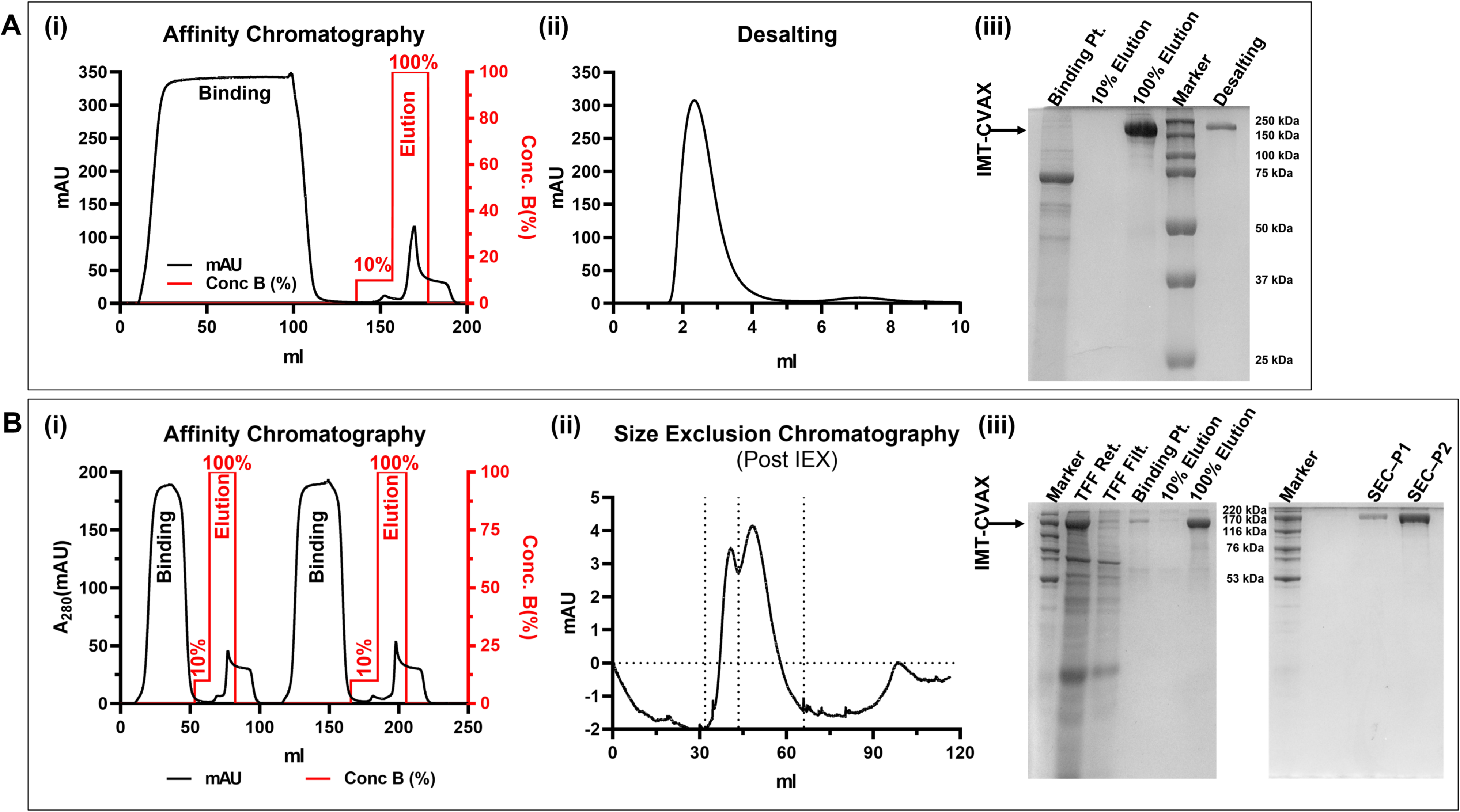
One-Step and Two-Step Purification of His-tagged IMT-CVAX: (A) Highly pure His-tagged IMT-CVAX was obtained after one round of affinity chromatography using affinity resin containing Cobalt. IMT-CVAX was eluted with 100% elution buffer after the column was thoroughly washed with 10% elution buffer. Buffer exchange was performed using a 5ml desalting column to bring the protein into its storage buffer. (i) is the IMAC chromatogram. After binding, the column was first washed with 10% elution buffer, and IMT-CVAX was subsequently eluted with 100% elution buffer. (ii) is the chromatogram generated while desalting. (iii) The SDS-PAGE of all the fractions collected during affinity chromatography and desalting. (B) If some additional bands were seen in the affinity-purified fraction, usually in the case where the viability of cells is <85% at the time of harvest, a second step of size-exclusion chromatography was performed to obtain highly pure IMT-CVAX. (i) is the chromatogram showing two back-to-back cycles of affinity chromatography. After binding, the column was first washed with 10% elution buffer, and IMT-CVAX was subsequently eluted with 100% elution buffer. (ii) is the size-exclusion chromatogram showing the two prominent peaks (P1 and P2) collected separately. (iii) The SDS-PAGE of all the fractions collected before and during affinity chromatography. A high Molecular Weight Protein Mixture (High Molecular Weight SDS Calibration Kit, GE Healthcare UK Limited, 17-0615-01/AB) was used as the molecular weight reference.

**SI Fig. 2:**
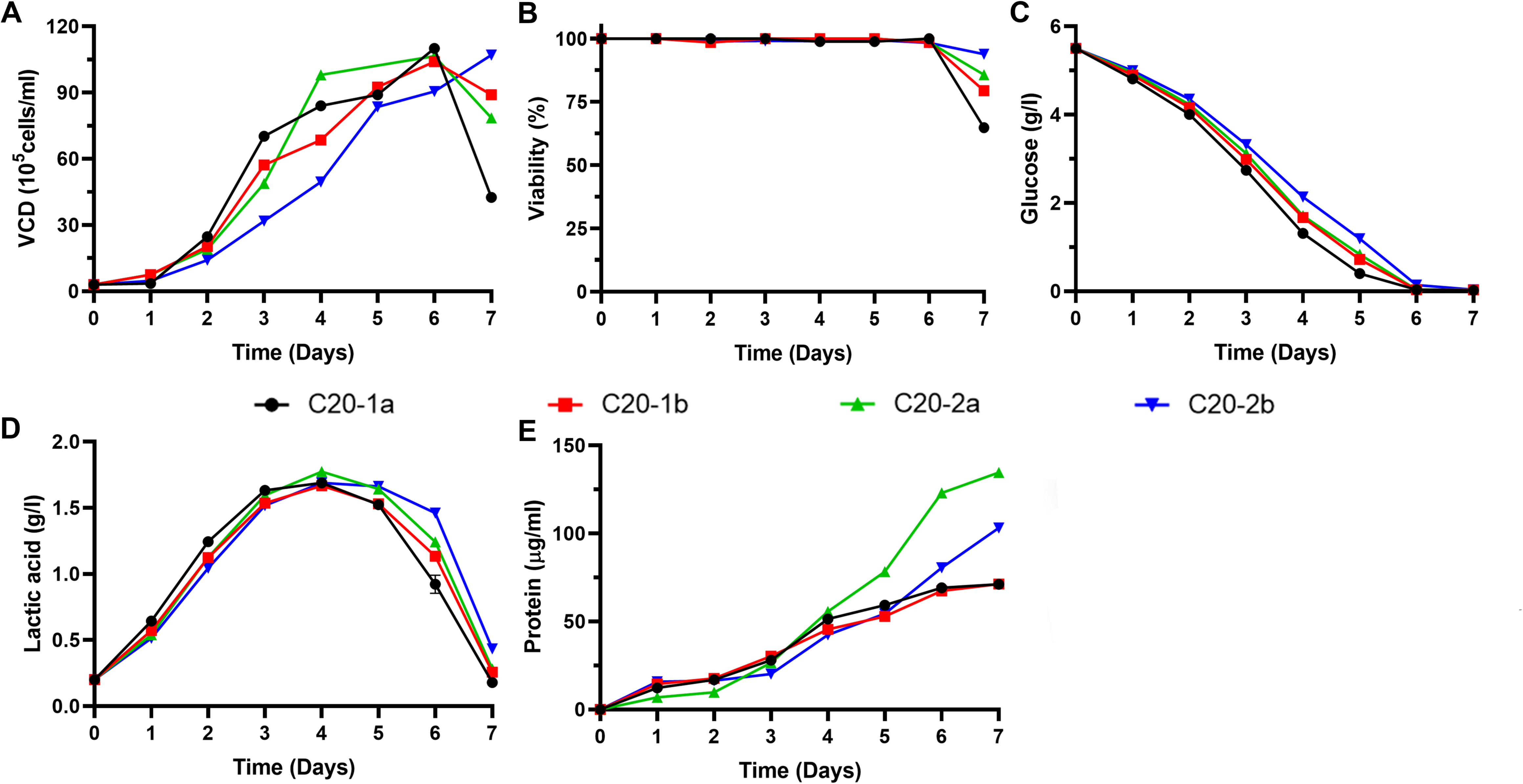
Volumetric Productivity of the Four Stable Cell Pools: Production phase of the 4 stable cell clones generated. C20-2a showed the highest expression of IMT-CVAX (e). The VCD (a) and Viability of C20-2b (b) at the time of harvest was the highest. (c) and (d) show glucose consumption and lactate production.

**SI Fig. 3:**
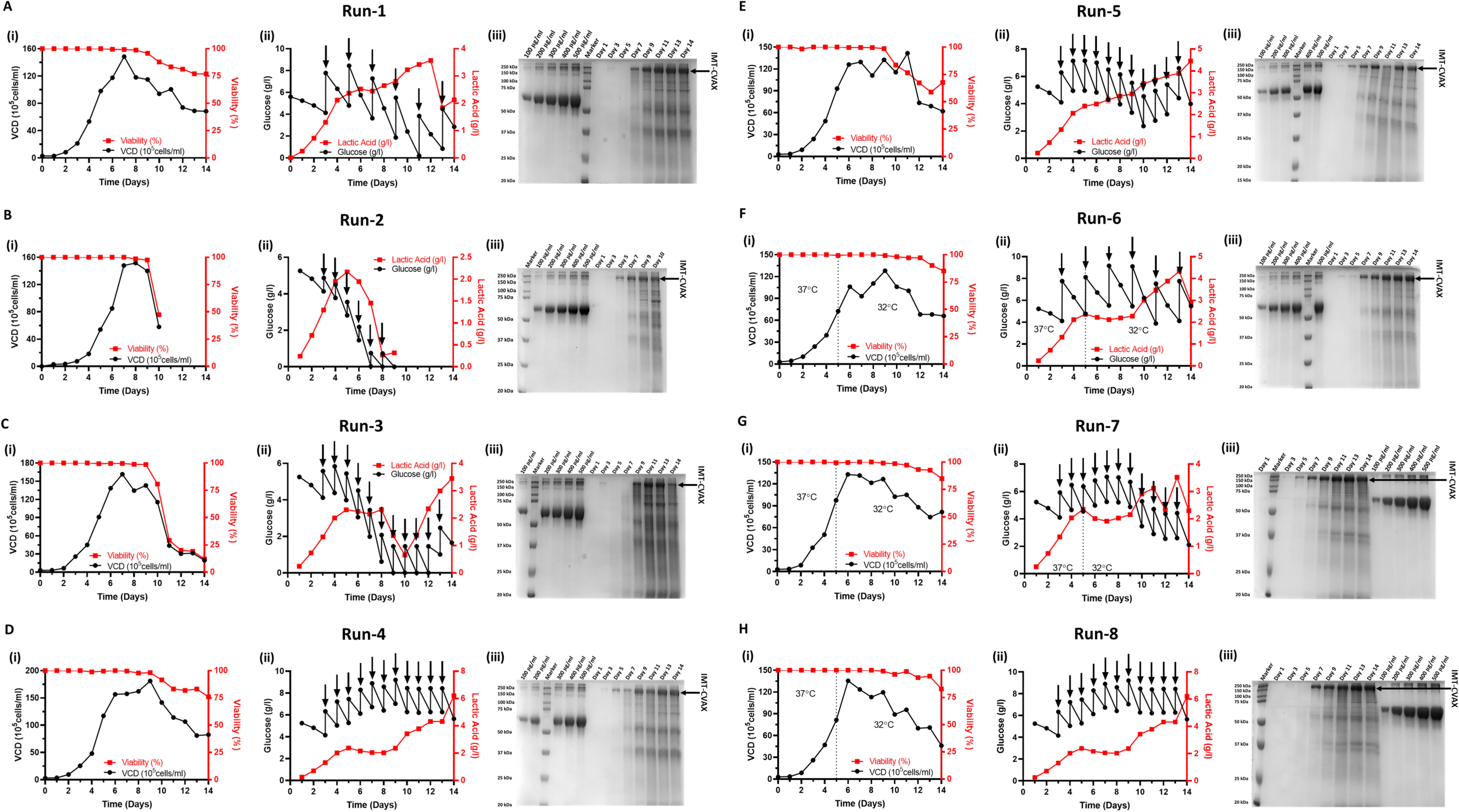
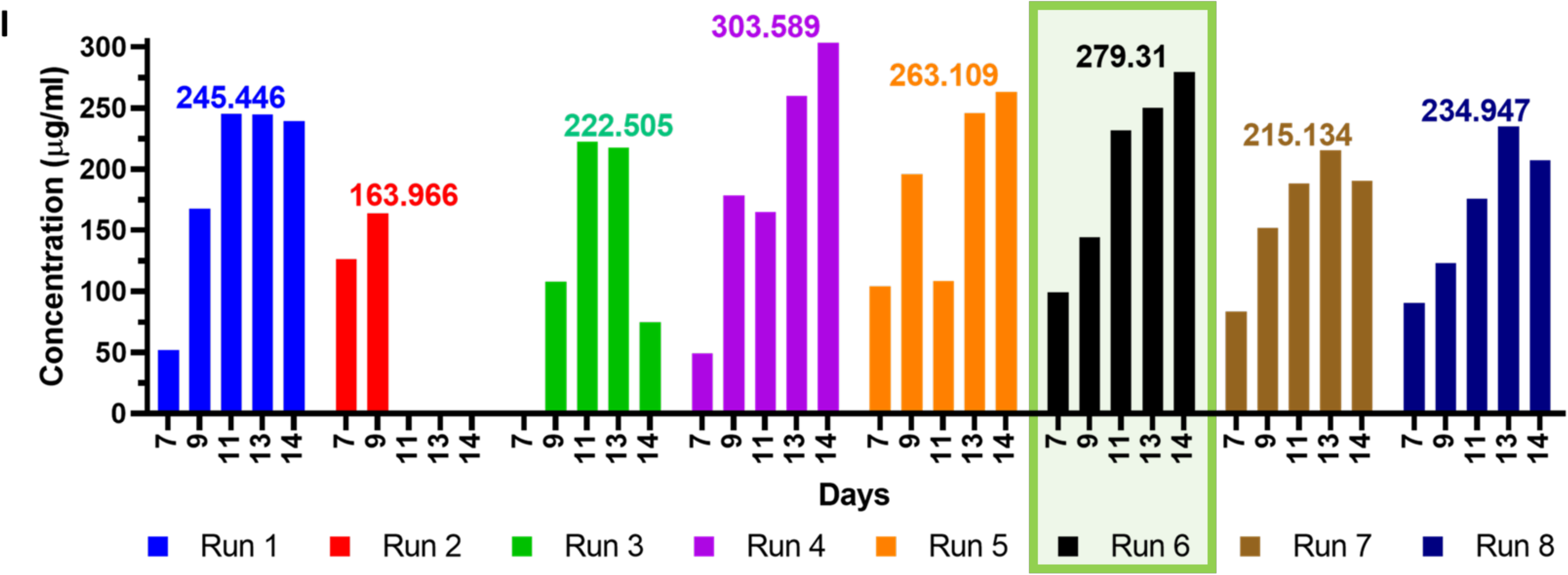
Optimization of Feed and Temp conditions: The high yields of IMT-CVAX in C20-2a were further increased under the feeding and temperature condition of Runs 4 (SI Fig.: D) and 6 (SI Fig.: F). In Runs 1-5, different feeding strategies were used. Runs 2 and 3 (1% and 2% daily feeding, respectively) experienced a drastic drop in viability after Day 9; hence were not considered for further optimizations. Runs 1, 4 and 5 maintained almost 70% viability till day 14. The feeding conditions of Runs 1,4, and 5 were mirrored in Runs 6-8 and additionally, the flasks were shifted from 37℃ to 32℃ on Day 5. The viabilities of these three flasks were maintained at >80% till day 14. Elevated lactic acid levels on day 14 were observed in Runs 7 and 8. Looking at the overall expression of IMT-CVAX, deduced by densitometric analyses (SI Fig.: I) of the SDS gels, Runs 4 and 6 showed the highest expression levels. Nevertheless, the viability of Run 6 was higher than that of Run 4, and, considering the ease of purification, it was deemed better and selected for bioreactor based expression. The feeding and temp conditions are mentioned in detail in Tables 1 and 2. IMT-CVAX concentrations of Runs 1-8, calculated using gel densitometry with known concentrations of BSA as reference. Run 6, with IMT-CVAX concentration of 279.31 µg/mL, was selected for upscaling to the Bioreactor level.

**SI Fig. 4:**
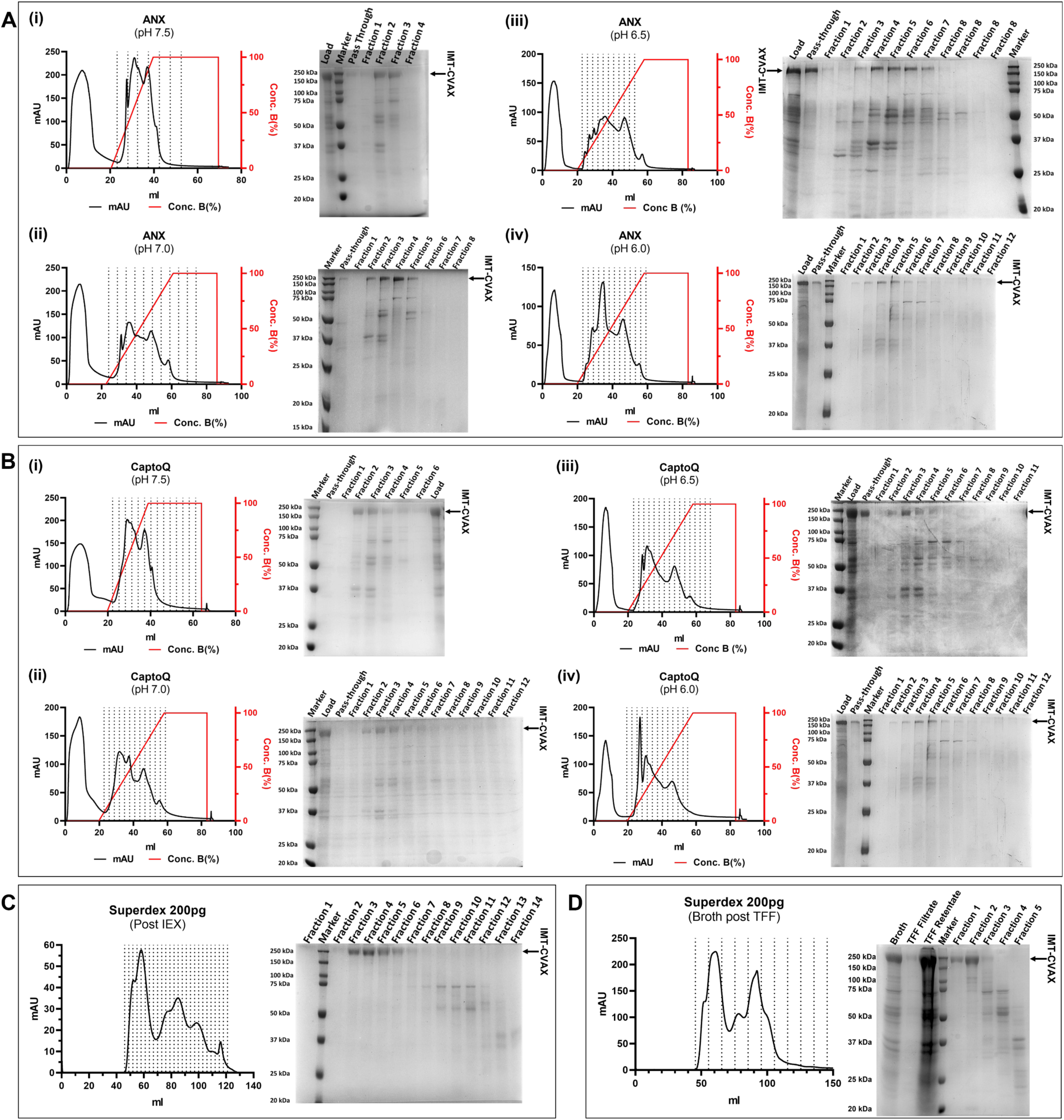
Optimization of IMT-CVAX Purification Process: Untagged IMT-CVAX was purified by anion-exchange chromatography. ANX (weak anion-exchange resin) (a) and CaptoQ (strong anion-exchange resin) (b) were tested at 4 different pH (7.5, 7.0, 6.5, and 6.0). The purest IMT-CVAX was obtained using CaptoQ along with buffers of pH 6.0. The recombinant spike protein was obtained as the pass-through, while the impurities bound on the column. A small fraction of IMT-CVAX did bind to the column. It was later eluted, pooled and then purified via size-exclusion chromatography using Superdex 200pg (c), which yielded highly pure IMT-CVAX. As a single-step chromatography, TFF retentate was directly loaded onto Superdex 200pg and, as is depicted in (d), pure IMT-CVAX was obtained in Fraction 1. However, fraction 2 had the majority of IMT-CVAX with traces of few other proteins impurities.

**SI Fig. 5:**
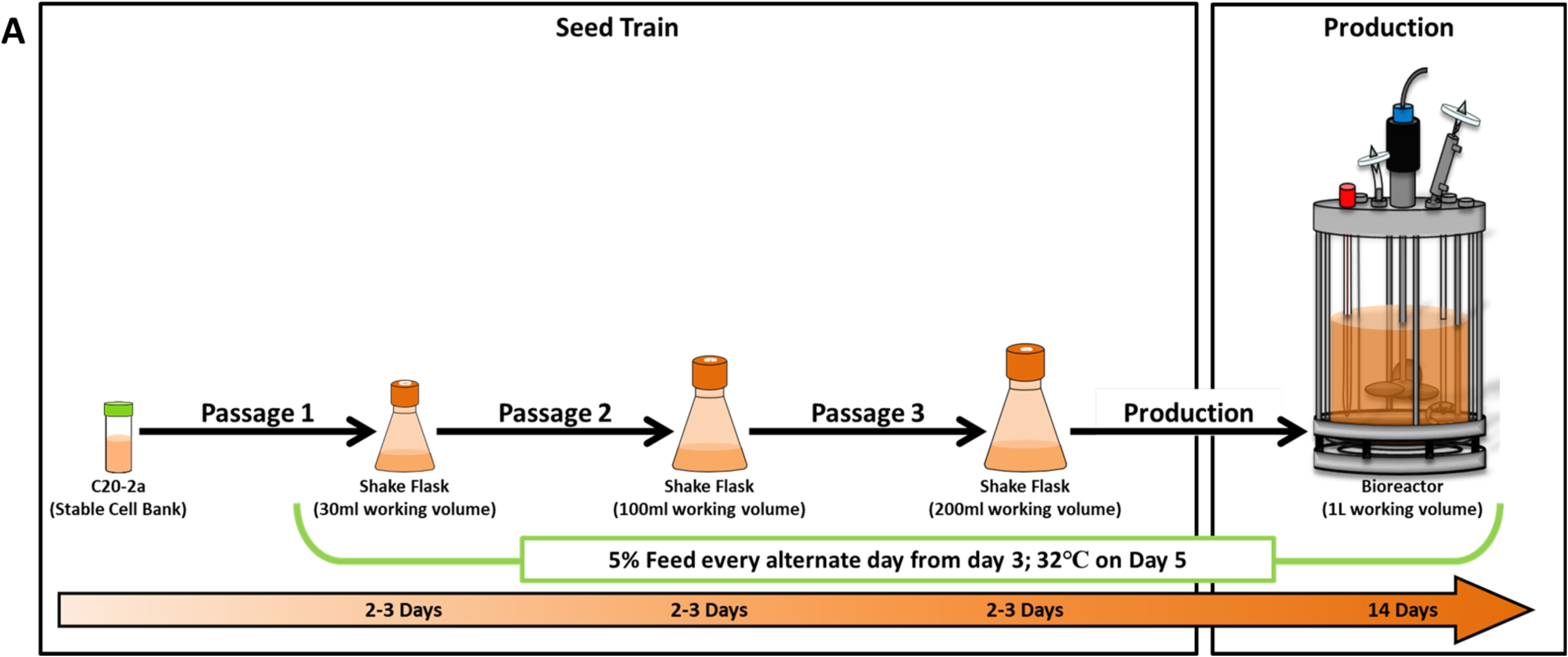
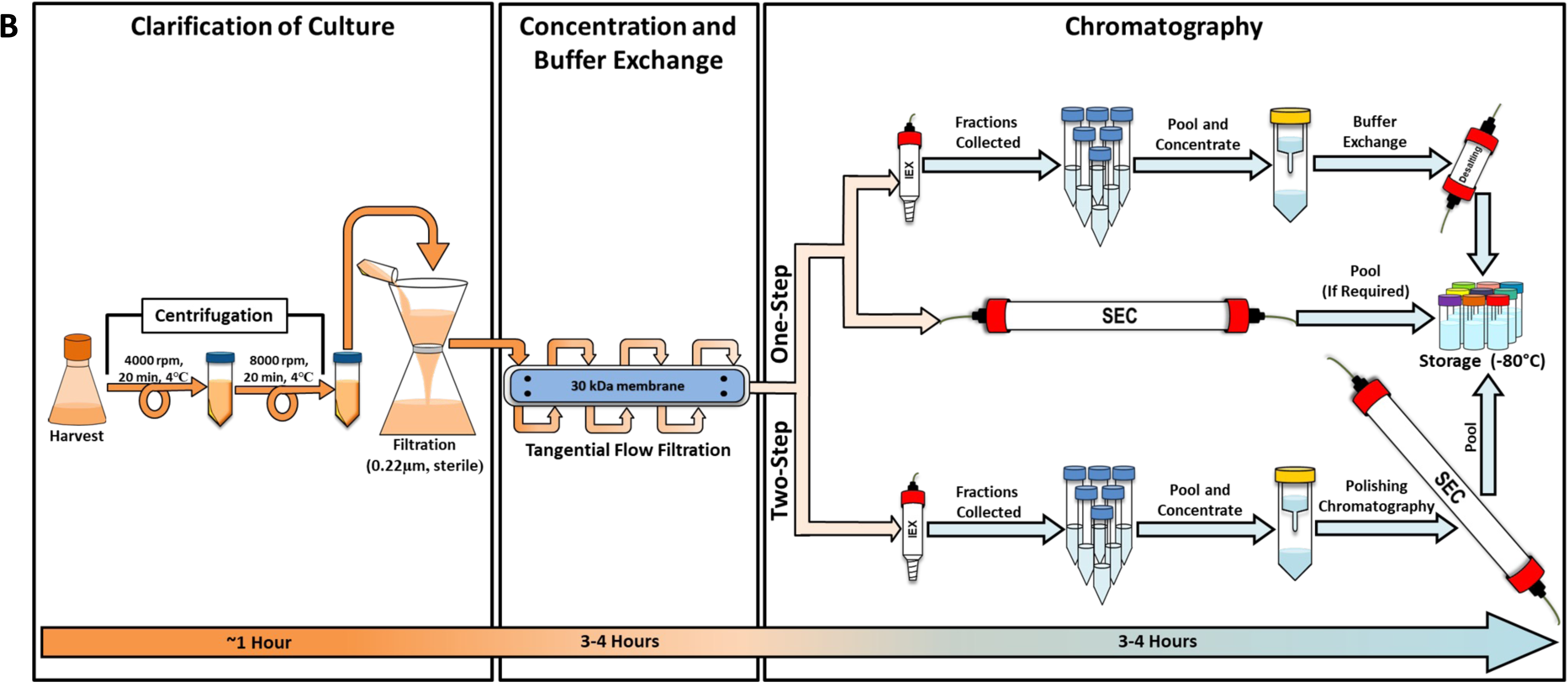
Overall process flow for expression and purification of IMT-CVAX: (A) Depicts the process flow of the selected stable cell pool, C20-2a, from flask to bioreactor level. The stock was first revived in 30ml media. The working volume of the culture was increased first to 100ml and then 200ml in passages 2 and 3, respectively. From the 200ml Flask, the culture was directly inoculated in the bioreactor with a working volume of 1L. The C20-2a batch was run for 14 days. The down-stream processing **(B)** involves first the separation of solids from the broth by centrifugation followed by filtration. The broth is then subjected to TFF where the spent media was exchanged with buffer, and smaller proteins are removed. TFF retentate is recovered and can be purified by either anion-exchange alone, or size-exclusion alone, or anion-exchange + size-exclusion (for viabilities <85% at the time of harvest). When performing anion-exchange alone, an additional desalting step is required, which replaces the elution buffer with the storage buffer, same as the size-exclusion buffer. Once purified, the fractions are pooled, concentrated (if required), aliquoted and stored at −80℃.

**Supplementary Table 1:**
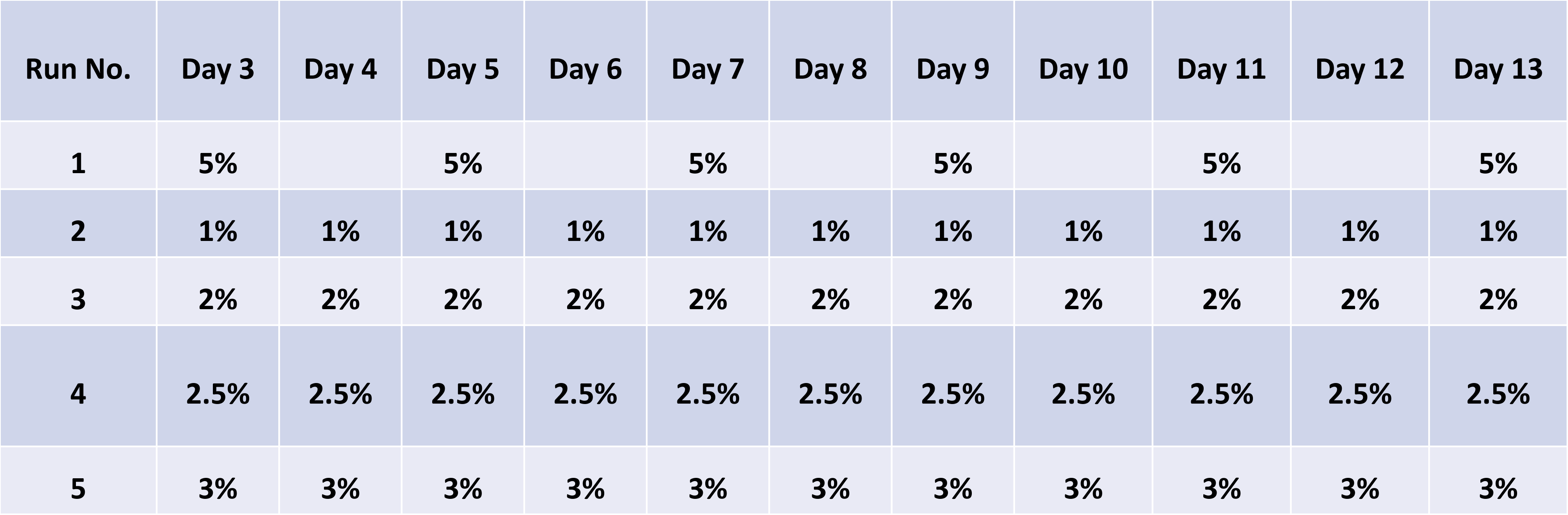
Feeding strategies used in the fed-batch experiments in which 37°C was kept constant for runs 1-5.

**Supplementary Table 2:**
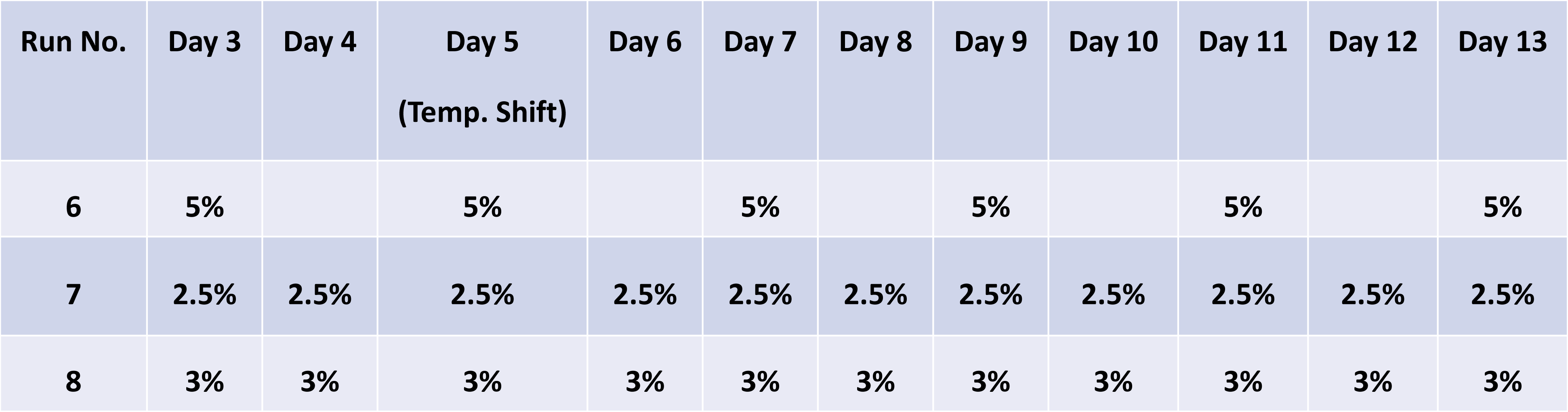
Feeding strategies used in the fed-batch experiments in which temperature shift from 37℃ to 32℃ was given from day 5 onwards.

